# PTEN neddylation aggravates CDK4/6 inhibitor resistance in breast cancer

**DOI:** 10.1101/2024.08.06.606911

**Authors:** Fan Liu, Weixiao Liu, Yawen Tan, Yaxin Shang, Sihui Ling, Xiaokun Jiang, Zhen Zhang, Shiyao Sun, Ping Xie

**Affiliations:** Department of Cell Biology, Laboratory for Clinical Medicine, Capital Medical University, Beijing 100069, China; Beijing lnstitute of Hepatology, Beijing Youan Hospital, Capital Medical University, Beijing 100069, China; Department of Breast and Thyroid Surgery, The Second People’s Hospital of Shenzhen, Shenzhen, Guangdong 518035, China; Department of Neurosurgery, Shandong Provincial Hospital affiliated to Shandong First Medical University, Jinan, Shandong 250021, China

## Abstract

The gradual emergence of a novel therapeutic approach lies in the restoration of tumor suppressive machinery. PTEN is a crucial negative regulator of the PI3K/Akt signaling pathway. Protein neddylation modification contributes to PTEN inactivation and fuels breast cancer progression. Here, we highlight that an elevated level of PTEN neddylation is markedly associated with resistance to palbociclib, a CDK4/6 inhibitor used in luminal subtype breast cancer patients. Mechanistically, PTEN neddylation activates the PI3K/Akt signaling pathway, and more notably, upregulates the activity of the AP-1 transcription factor. Our data showed that PTEN neddylation stabilizes JUND, a transcription factor involved in the AP-1 complex, by disrupting its interaction with the E3 ubiquitin ligase ITCH. Consequently, activated JUND leads to the release of cytokines and chemokines, which in turn may drive an inflammatory tumor microenvironment, potentially contributing to drug resistance. Then, we identified Echinacoside as a potent inhibitor of PTEN neddylation both *in vivo* and *in vitro* by disrupting its interaction with XIAP, the E3 ligase responsible for PTEN neddylation. Combination of Echinacoside effectively overcome resistance to palbociclib in breast cancer treatment. These findings highlight targeting PTEN neddylation as a promising strategy for restoring tumor suppressor activity and overcoming resistance.

## INTRODUCTION

Breast cancer is a heterogeneous disease with multiple intrinsic tumor subtypes. The luminal breast cancer subtype, characterized by the expression of hormone receptor-positive (HR+) and human epidermal growth factor receptor 2-negative (HER2-) accounts for 70 % of breast cancer patients [1, 2]. Cyclin D-dependent Kinases 4 and 6 inhibitors (CDK4/6i) have been approved by the FDA for the treatment of luminal breast cancer [3–5]. Although patients with advanced disease have benefited from CDK4/6 inhibitors, drug resistance still remains a significant clinical challenge [6–8]. Many factors contribute to CDK4/6i resistance, including CDK4/6 and Cyclin D1 overexpression, p16 amplification, loss of p53, loss of retinoblastoma (Rb) and over-activation of CDK2. Notably, resistance to CDK4/6 inhibitors is also prominently characterized by over-activated PI3K/Akt signaling [9–13]. However, although PI3K inhibitors could prevent resistance to CDK4/6 inhibitors, they failed to re-sensitize cells once resistance had been acquired [14, 15]. PTEN (phosphatase and tension homologue deleted from chromosome 10) is a well-known negative regulator of the PI3K signaling pathway and governing fundamental cellular processes [16–19]. Recently, PTEN loss is considered as a bona fide mechanism of resistance to CDK4/6 inhibitors. Loss of PTEN increases PI3K/Akt activity and increases activity of CDK4/6 [20–22]. Therefore, targeting the restoration of PTEN tumor suppressor activity is believed to be a more effective approach in tumor treatment [23, 24].

Nedd8, a ubiquitin-like protein covalently conjugated to substrates similar to ubiquitin, mediates protein post-translational modification called neddylation [25]. Neddylation involves the covalent attachment of Nedd8 to a substrate through an ATP-dependent cascade involving the Nedd8-activating enzyme (E1, UBA3/APPBP1), Nedd8-conjugating enzyme (E2, UBE2M and UBE2F), and Nedd8 ligases (E3). The substrates of neddylation include both cullin proteins and non-cullin proteins [26, 27]. Over-active neddylation modification has been indicated to be closely correlated with cancer development. The first-in-class neddylation E1 inhibitor, MLN4924 (pevonedistat), has entered in several phases of clinical trials for tumor therapy [28]. Additionally, several inhibitors targeting E2s have shown well anti-tumor activity in mice models [29]. Up to now, none of them is granted by FDA for the treatment of cancer. It’s still necessary to discover more potent and selective neddylation inhibitors for clinical trials. Our previous study found that high levels of PTEN neddylation, which lead to PTEN inactivation, enhance the activity of the PI3K/Akt pathway, promote PTEN nuclear import, and accelerate breast tumor proliferation and invasion [30, 31]. Here, we found that PTEN neddylation is elevated in luminal-type breast cancer patients with resistance to palbociclib by promoting JUND protein stability and AP-1 transcriptional activation. Furthermore, we proposed that Echinacoside (ECH), a natural phenylethanoid glycoside in *Echinacea angustifolia* DC, acts as a potent PTEN neddylation inhibitor. Combined treatment of ECH therapeutically blunted palbociclib resistance and enhanced drug sensitivity. Therefore, we provide an innovative therapeutic strategy to limit cancer drug resistance through PTEN reactivation by blocking neddylation modification.

## RESULTS

### Resistance to CDK4/6 inhibitor upregulates PTEN neddylation in breast cancer

We previously found that elevated PTEN neddylation in breast cancer correlates with poor clinical outcomes [30], and we confirmed those results here (Fig. 1A, B). We observed that PTEN neddylation was upregulated in all breast cancer subtypes, particularly in luminal-type cancers, while PTEN was downregulated in all subtypes without distinction (Fig. 1C). Detailed pathological information was showed in Supplementary Table 1. Palbociclib, one kind of CDK4/6 inhibitor, has been widely used for the treatment of advanced breast cancer patients. Using the antibody that specifically recognized PTEN neddylation, our results showed that PTEN neddylation was markedly upregulated in palbociclib-resistant breast cancer patients with recurrence *in situ* and liver metastasis, while PTEN remained lowly expressed in drug-resistant patients (Fig. 1D). Pathological data of the patients were provided (Supplementary Fig. 1A). Next, we monitored MMTV-*PyMT* (C57BL/6J background) transgenic mice, which spontaneously develop luminal breast tumors, as a model for late-stage luminal breast cancer [32–34]. Following spontaneous tumor formation, breast tumors exhibited a significant reduction in volume and weight after about 20 days of palbociclib treatment, suggesting that this period corresponded to the drug-sensitive phase of palbociclib. Notably, as the duration of continuous treatment was extended, the tumor volumes and weights ceased to decline further, indicating the emergence of drug resistance. This drug resistance could persist throughout the 50-day treatment period (Fig. 1E-H). Moreover, compared with the low-dose group (75 mg/kg) that developed drug resistance at 20 days, the high-dose group (100 mg/kg) showed that the tumor volume and weight no longer continued to decrease after 25 days, indicating the onset of the drug resistance. Palbociclib-resistant tumors showed no significant difference in size or weight compared to controls, and were marked larger than in the sensitive group at 35-day (Fig. 1I-L).

**Fig. 1.**
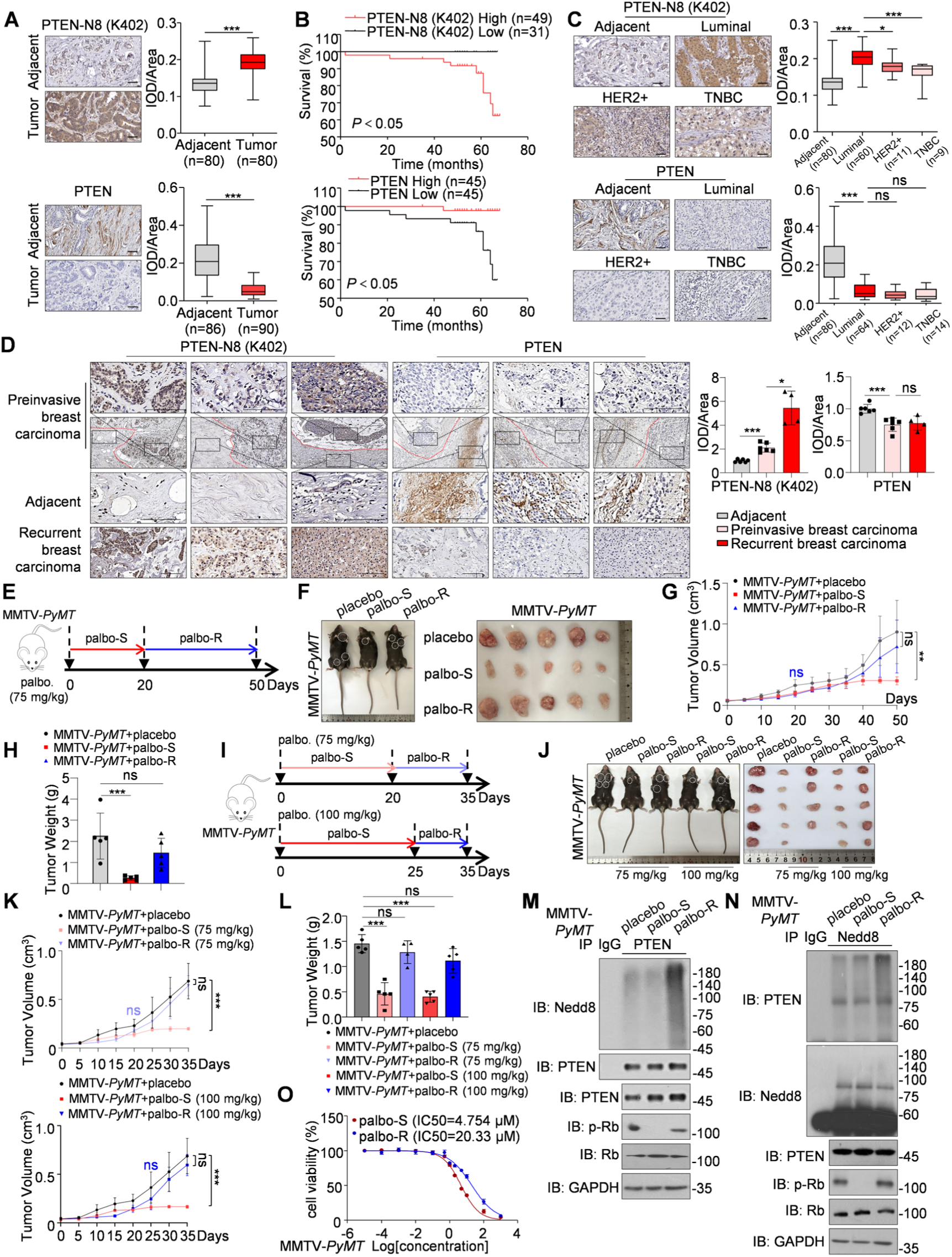
Palbociclib-resistant breast cancer cells express higher levels of PTEN neddylation modification. **A** Representative image from immunohistochemical staining of PTEN and PTEN neddylation on K402 site in breast cancer and matched adjacent tissues. ImageJ was used to perform Semi-quantitative analysis. Data were analyzed using Student’s *t*-test. Scale bars, 50 µm. PTEN and PTEN neddylation on K402 expression are shown as box plots. **B** Kaplan-Meier plot of overall survival of patients with breast carcinomas. A log-rank test was used to show differences between groups (red indicates high PTEN or PTEN-N8 K402 expression; black indicates low PTEN or PTEN-N8 K402 expression). **C** Representative images from immunohistochemical staining of PTEN and neddylated PTEN on K402 site in different breast cancer subtypes and adjacent tissues are shown. Scale bars, 50 µm. Data were analyzed using the one-way ANOVA test. PTEN and PTEN neddylation on K402 expression are shown as box plots. **D** Representative images from immunohistochemical staining of PTEN and PTEN neddylation on K402 in preinvasive breast cancer tissues (n=6), palbociclib resistance breast cancer tissues (n=6) and adjacent tissues (n=4). Data were analyzed using Student’s *t*-test. Scale bars, 100 µm. The protein level of PTEN and PTEN neddylation on K402 expression are shown as bar plots. **E-H** To establish the palbociclib-resistant mouse model, the palbociclib-resistant MMTV-*PyMT* mouse model was established by palbociclib (75 mg/kg) gavage daily for 50 days and the palbociclib-sensitive group were administered palbociclib (75 mg/kg) by gavage daily for 20 days. The three conditions were compared in the same time (n=5) (**E**). The breast tumors were removed and photographed (**F**). Tumors were isolated, volumes (**G**) and their weights (**H**) were measured. Data are presented as means ± S.D. **I-L** To establish the palbociclib-resistant mouse model, the palbociclib-resistant MMTV-*PyMT* mouse model was established by palbociclib (75 mg/kg or 100 mg/kg) gavage daily for 35 days and the palbociclib-sensitive group were administered palbociclib (75 mg/kg or 100 mg/kg) by gavage daily for 20 days (placebo group n=5; palbo-S-75mg/kg group n=5; palbo-R-75mg/kg group n=4; palbo-S-100mg/kg group n=5; palbo-R-100mg/kg group n=5). The three conditions were compared in the same time (**I**). The breast tumors were removed and photographed (**J**). Tumors were isolated, volumes (**K**) and their weights (**L**) were measured. Data are presented as means ± S.D. **M** Immunoblot analysis of anti-PTEN immunoprecipitate and cell lysates derived from spontaneous breast tumors of MMTV-*PyMT* mice in the indicated three groups. **N** Immunoblot analysis of anti-Nedd8 immunoprecipitate and cell lysates derived from spontaneous breast tumors of MMTV-*PyMT* mice in the indicated three groups. **O** IC50 of palbociclib-resistant breast tumor cells obtained from MMTV-*PyMT* mice (MMTV-*PyMT*+palbo-S and MMTV-*PyMT*+palbo-R). The cells were seeded in 96-well plates treated with increasing concentration (0.01 nM-1000 µM) of palbociclib and then analyzed for CCK8 assay. *P* values were calculated by Student’s *t*-test. (**A**), log-rank test (**B**) and one-way ANOVA test (**C**, **D**, **G**, **H**, **K**, **L**). Error bars, ± S.D. ns, not significant, \**P* < 0.05, \*\**P* < 0.01, \*\*\**P* < 0.001.

Rb is a tumor suppressor that is phosphorylated during the transition from G1 to S phase of the cell cycle by CDK4/6. When CDK4/6 inhibitors are ineffective, phosphorylation of the Rb protein would become uncontrolled [35]. Therefore, the increased phosphorylation of the Rb protein in the mice also indicated the occurrence of drug resistance (Fig. 1M, N input group; Supplementary Fig. 1B). More importantly, high levels of PTEN neddylation were detected in palbociclib-resistant breast tumor tissues of MMTV-*PyMT* mice, while the expression of PTEN remained no obviously changed (Fig. 1M, N and Supplementary Fig. 1B). Additionally, we found that the IC50 of breast tumor cells isolated from palbociclib-resistant MMTV-*PyMT* mice was significantly higher compared to the palbociclib-sensitive group (Fig. 1O), indicating the development of drug resistance. Taken together, our data revealed that PTEN neddylation exhibited elevated levels in palbociclib-resistant breast cancer.

### Increased XIAP leads to enhanced PTEN neddylation in palbociclib-resistant breast cancer

To explore the underlying mechanisms of elevated PTEN neddylation in palbociclib-resistant breast cancer, we generated acquired palbociclib-resistant breast cancer stable cell lines (MCF-7-PR). Dose-response curves results showed that palbociclib-resistant cells were more resistant to palbociclib than the parental cells (Fig. 2A). Compared with the parental cells, the palbociclib-resistant cells exhibited accelerated G1/S phase and cell proliferation ability after treatment of palbociclib (Fig. 2B, C). Consistently, PTEN neddylation was upregulated in acquired palbociclib-resistant breast cancer cells (Fig. 2D). Moreover, the ubiquitination of PTEN was unchanged in palbociclib-resistant cells (Supplementary Fig. 1C). Compared to other drug-resistant breast cancer cell lines, such as MCF-7 sublines resistant to doxorubicin (MCF-7/ADR), cisplatin (MCF-7/DDP), and paclitaxel (MCF-7/PTX), PTEN neddylation was specifically increased in palbociclib-resistant breast cancer cells (Fig. 2E). Short-term drug treatment could induce adaptive resistance in cells [13]. We noticed that the phosphorylation of Rb, initially reduced by palbociclib treatment, recovered after prolonged drug exposure, which indicated adaptive resistance was occurred (Fig. 2F). Under the adaptive resistance, PTEN neddylation was increased and could be reduced by MLN4924, which was the inhibitor of neddylation E1 (Fig. 2F, G left panels). For ER negative cells, such as glioma cells T98G, show reduced palbociclib sensitivity [36]. As expected, no obvious changes in PTEN neddylation or p-Rb were detected when the T98G cells were treated with palbociclib (Fig. 2G right panels).

**Fig. 2.**
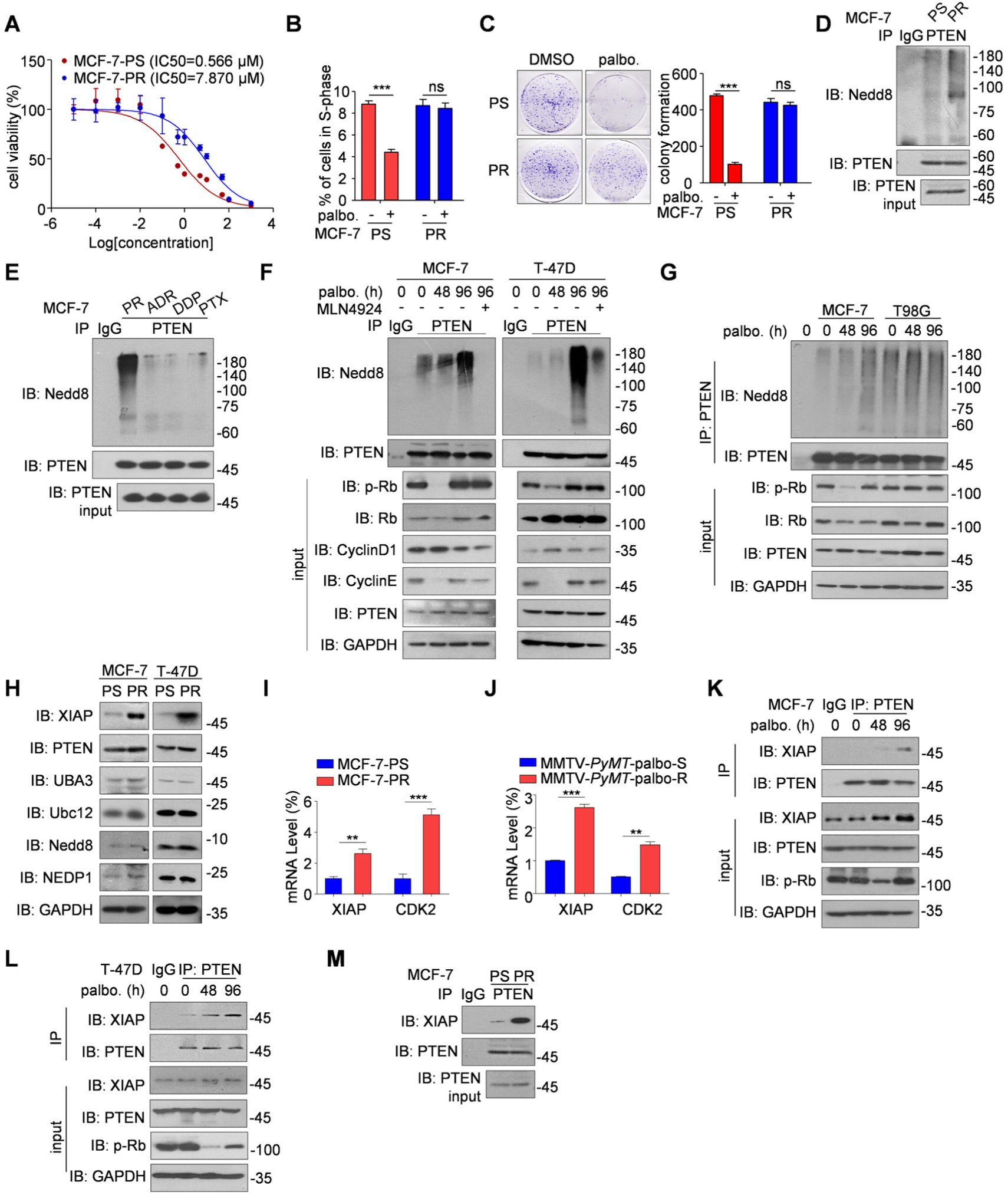
Upregulation of XIAP increases the PTEN neddylation in palbociclib-resistant breast cancer. **A** IC50 of MCF-7-PS (palbociclib sensitivity, PS) and MCF-7-PR (palbociclib resistance, PR) cells. MCF-7-PR cells and MCF-7-PS cells were seeded in 96-well plates and then treated with increasing concentrations (0.01 nM-1000 µM) of palbociclib and then analyzed for CCK8 assay. **B** MCF-7-PS and MCF-7-PR cells are treated with palbociclib (0.5 µM, 24 h) or not. The percentage of cells in G1/S-phase is depicted. Data are presented as the means ± S.D. (n=3). **C** Colony formation assay was performed in the MCF-7-PS cells and MCF-7-PR cells and the cells were treated with palbociclib (0.1 µM, 15 days). Data are presented as means ± S.D. Results are from a representative experiment performed in triplicate. ImageJ was used to perform quantitative analysis. **D** Immunoblot analysis of anti-PTEN immunoprecipitate from MCF-7-PS and MCF-7-PR cells to detect PTEN neddylation *in vivo*. **E** Immunoblot analysis of anti-PTEN immunoprecipitate from MCF-7 drug resistant cells including MCF-7-PR, MCF-7 sublines resistant to doxorubicin (MCF-7-ADR), cisplatin (MCF-7-DDP) and paclitaxel (MCF-7-PTX) cells. **F** MCF-7 cells and T-47D cells were treated with palbociclib (0.5 μM, 0 h/48 h/96 h). The right panel was treated with palbociclib (0.5 μM, 96 h) and MLN4924 (1 μM, 12 h) simultaneously. The cells were collected and lysed for immunoblot analysis with indicated antibodies. **G** Immunoblot analysis of anti-PTEN immunoprecipitate from MCF-7 cells and T98G cells. MCF-7 cells and T98G cells were respectively treated with palbociclib (0.5 μM, 0 h/48 h/96 h). **H** Western blot analysis of XIAP, PTEN, UBA3, Ubc12, Nedd8, NEDP1 in palbociclib sensitivity and resistance MCF-7 and T-47D cells. **I** Analysis of CDK2 and XIAP mRNA levels in MCF-7-PS/PR cells. **J** Analysis of CDK2 and XIAP mRNA levels in MMTV-*PyMT*-palbo-S/palbo-R tissues. **K, L** Immunoblot analysis of anti-PTEN immunoprecipitate and WCL from adaptative palbociclib resistance in MCF-7 (**K**) and T-47D (**L**). **M** Coimmunoprecipitation of XIAP after immunoprecipitation of the PTEN protein in palbociclib sensitivity and resistance MCF-7 cells. *P* values were calculated by one-way ANOVA test (**B, C**) and Student’s *t*-test (**I, J**). Error bars, ± S.D. ns, not significant, *\*\*P* < 0.01, *\*\*\*P* < 0.001.

Then, we intended to explore the mechanism of high PTEN neddylation in palbociclib-resistant breast cancer. Notably, our data revealed that XIAP, the E3 ligase for PTEN neddylation [30], was upregulated in palbociclib-resistant breast cancer cells, while no significant changes were observed in Nedd8 E1-UBA3, Nedd8 E2-Ubc12, and Nedd8 (Fig. 2H). Consistent with previous studies showing compensatory CDK2 activation upon resistance elevates XIAP mRNA levels [37, 38], we observed increased XIAP and CDK2 mRNA in both drug-resistant cells and mouse models (Fig. 2I, 2J). The increased XIAP enhanced the interaction with PTEN in both early adaptative palbociclib-resistant and acquired palbociclib-resistant breast cancer cells (Fig. 2K-M). The above data suggested that high PTEN neddylation in palbociclib-resistant breast cancer cells was likely due to XIAP upregulation and increased XIAP-PTEN interaction.

### PTEN neddylation enhances the protein stability of JUND

In previous studies, we have demonstrated that Nedd8 is covalently attached to lysine residues at positions 197 and 402 of PTEN [30]. To examine the role of PTEN neddylation on CDK4/6i resistance, we performed RNA-seq in palbociclib-resistant breast cancer cells (Fig. 3A) and PTEN neddylation deficient cells (2KR, K197 and K402 sites mutated to R) (Fig. 3B). Other than the PI3K/Akt signaling pathway, which has been believed to be highly associated with CDK4/6i resistance and PTEN neddylation, we noticed that MAPK, as well as JAK/STAT signaling pathways were also enriched (Fig. 3A, B). GSEA analysis further indicated that MAPK signaling pathway obviously was also enriched in both palbociclib-resistant breast cancer cells (Fig. 3C) and PTEN neddylation deficient cells (Fig. 3D). Similar data were shown in the related dataset (GSE222367) (Supplementary Fig. 2A). AP-1 (Activator Protein-1) transcription factors are heterodimers or homodimers composed of JUN and FOS proteins. AP-1 is known to be activated by MAPK signaling pathways through the phosphorylation of JUN and FOS proteins, and in turn, AP-1 also regulates the expression of genes involved in MAPK signaling pathways [39]. The reporter gene assays revealed that AP-1 transcription factor activity was significantly increased after palbociclib resistance developed (Fig. 3E). Further heat map data showed that palbociclib resistance led to the upregulation of AP-1 related transcription factors, including *FOSB*, *FOSL*, *c-FOS*, *c-JUN*, *JUND*, and *JUNB* (Fig. 3F and Supplementary Fig. 2B). Conversely, PTEN neddylation deficiency decreased the expression of those genes (Fig. 3G). Then, we detected the protein levels of the AP-1 transcriptional factors and noticed that there was a markable increase of JUND in palbociclib-resistant cells (Fig. 3H). Meanwhile, the activated form of JUND, phosphorylated JUND, was also upregulated when cells were resistant to palbociclib (Fig. 3I). Similar results were also found in early adaptive palbociclib-resistant MCF-7 cells and palbociclib-resistant breast cancer tissues in MMTV-*PyMT* mice, but not in ER-negative T98G cells (Fig. 3J, K). Next, analysis of nuclear and cytoplasmic fractions revealed that, in palbociclib-resistant breast cancer cells, JUND and phosphorylated JUND were significantly elevated, primarily in the nucleus (Fig. 3L). Finally, the results showed that JUND was also markedly upregulated in palbociclib-resistant breast cancer patients (Fig. 3M). These results strongly suggest that the MAPK/AP-1 signaling pathway is a key mechanism of resistance to palbociclib, and PTEN neddylation upregulates JUND protein expression while regulating AP-1 activity.

**Fig. 3.**
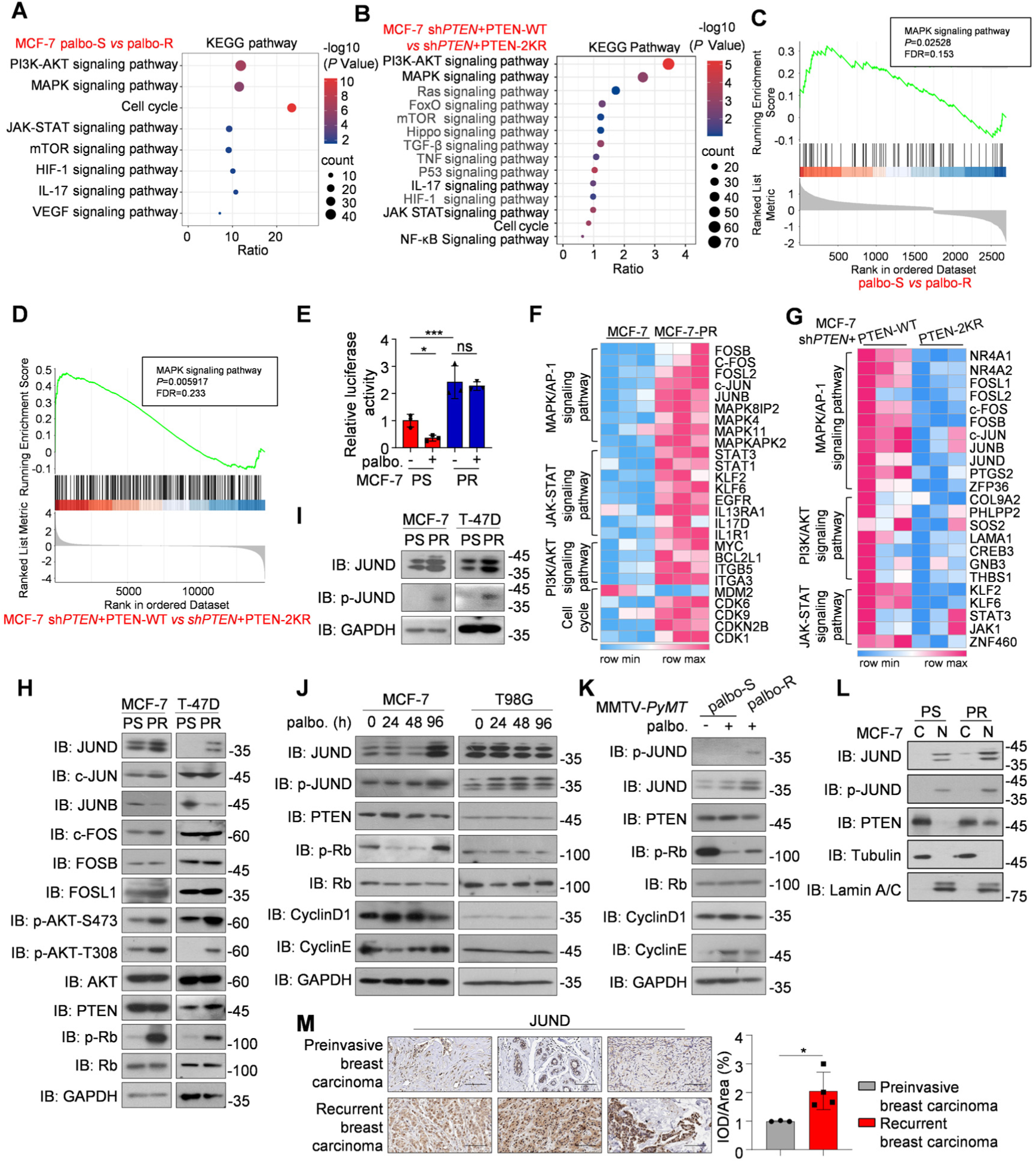
PTEN neddylation enhances JUND protein stability. **A, B** The bubble charts depict the top ranked pathway analyzed from the KEGG pathway database. Enriched KEGG pathways were identified, which included upregulated or downregulated genes modulated by the palbociclib resistance MCF-7 cells (**A**) and MCF-7 cells harboring a PTEN neddylation mutation (**B**). **C, D** GSEA plots of the palbociclib resistance MCF-7 cells (**C**) and PTEN neddylation deficiency MCF-7 cells (**D**). **E** AP-1 luciferase reporter plasmid was transfected in the palbociclib sensitivity and resistance MCF-7 cells. Reporter activity was assayed and represented as the means ± S.D. of three separate experiments. **F, G** Heat map of the palbociclib resistance MCF-7 cells (**F**) and PTEN neddylation deficiency MCF-7 cells (**G**). **H, I** Immunoblot of WCL from acquired palbociclib-sensitive or palbociclib-resistant cells. **J** Western blot analysis of lysates from MCF-7 cells treated for 24 to 96 hours with palbociclib (Palbo). Addition of fresh vehicle or drug every 24 hours over a 96-hour period. **K** The cells obtained from breast tumors of MMTV-*PyMT* mice were collected and lysed for immunoblot analysis. **L** Immunoblot of Nuclear (N) *vs* Cytoplasmic (C) fractionation from palbociclib sensitivity and resistance MCF-7 cells. **M** Representative images from immunohistochemical staining of JUND in palbociclib-resistant breast tumors (n=4) and matched adjacent tissues (n=3). ImageJ was used to perform Semi-quantitative analysis. Scale bars, 50 µm. *P* values were calculated by Student’s *t*-test. Error bars, ± S.D. ns, not significant, \**P* < 0.05, \*\**P* < 0.01, \*\*\**P* < 0.001.

### PTEN neddylation decreases JUND ubiquitination

Next, we intended to explore the mechanism by which PTEN neddylation promotes JUND protein expression, IP analysis indicated that the ubiquitination of JUND was obviously decreased in palbociclib-resistant breast cancer cells (Fig. 4A, B). We generated a fusion protein (PTEN-Nedd8) by linking Nedd8 to the C-terminus of PTEN to mimic neddylated PTEN. Neddylation would promote PTEN nuclear translocation. Therefore, a nuclear localization sequence (NLS) was fused to the N-terminus of PTEN to generate NLS-PTEN for comparison [30, 40, 41]. Then we observed an increased expression level of JUND and phosphorylated JUND in sh*PTEN*+PTEN-Nedd8 MCF-7 cells, but the opposite effect was found in sh*PTEN*+PTEN-NLS MCF-7 cells (Fig. 4C). Compared with NLS-PTEN, JUND showed a higher affinity for neddylated PTEN (Fig. 4D). Moreover, enhanced interaction between neddylated PTEN and JUND reduced JUND ubiquitination (Fig. 4E). Next, we confirmed that JUND expression was significantly increased in tumor tissues compared with matched adjacent normal tissues, particularly in luminal-type breast cancer (Supplementary Fig. 2C, D). By comparison, the levels of JUND and neddylated PTEN were positively correlated in luminal-type breast cancer (Supplementary Fig. 2E, F). Detailed pathological information was in Supplementary Table 2.

**Fig. 4.**
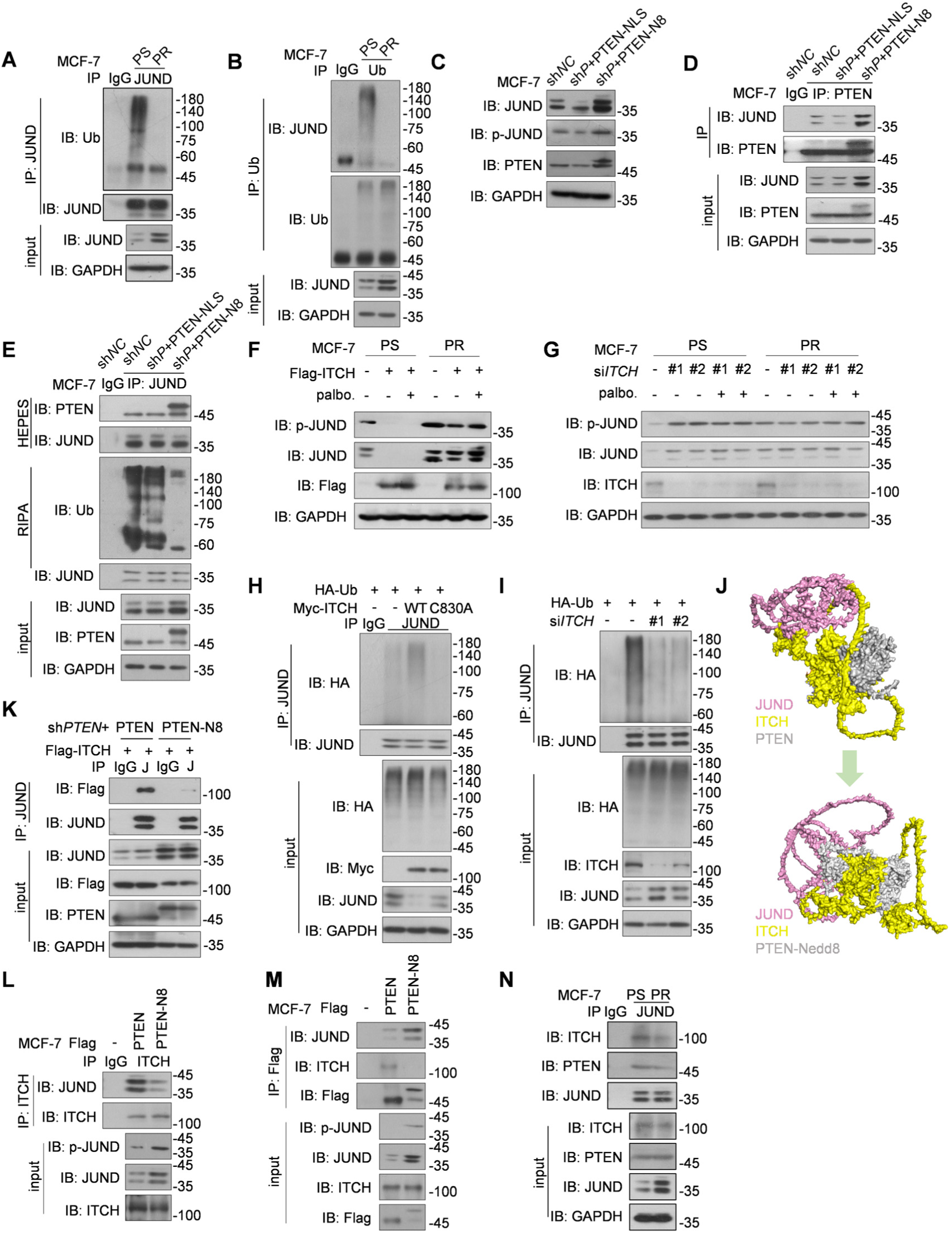
PTEN neddylation hinders the ITCH-mediated ubiquitination degradation of JUND. **A** Immunoprecipitation of JUND in the MCF-7-PS and MCF-7-PR cells and analysis using immunoblotting with the indicated antibodies. **B** Immunoprecipitation of Ub in the MCF-7-PS and MCF-7-PR cells and analysis using immunoblotting with the indicated antibodies. **C** Immunoblot of JUND and p-JUND in the MCF-7 sh*NC*, MCF-7 sh*PTEN*+PTEN-NLS and MCF-7 sh*PTEN*+PTEN-N8 cells. **D, E** Immunoprecipitation of PTEN (**D**) and JUND (**E**) in the MCF-7 sh*NC*, MCF-7 sh*PTEN*+PTEN-NLS and MCF-7 sh*PTEN*+PTEN-N8 cells. **F** Flag tagged-ITCH was transfected into the MCF-7-PS cells and MCF-7-PR cells and immunoblotted with the indicated antibodies. The cells were treated with palbociclib (0.5 μM for 48 h). **G** Si*ITCH* was transfected into the MCF-7-PS cells and MCF-7-PR cells and immunoblotted with the indicated antibodies. The cells were treated with palbociclib (0.5 μM for 48 h). **H** Immunoprecipitation of JUND in HEK293T cells transfected with plasmids of HA-tagged-Ub, Myc-tagged-ITCH and Myc-tagged-ITCH C830A and analysis using immunoblot analysis with the indicated antibodies. **I** Immunoprecipitation of JUND in HEK293T cells transfected with si*ITCH* and analysis using immunoblot analysis with the indicated antibodies. **J** Structure simulation of ITCH (yellow), JUND (pink) and PTEN or PTEN-Nedd8 (grey) by AlphaFold2.0. **K** MCF-7 sh*PTEN*+PTEN-WT and MCF-7 sh*PTEN*+PTEN-N8 cells transfected with plasmids for Flag-tagged ITCH and analyzed using immunoblot analysis with the indicated antibodies. **L, M** Flag tagged-PTEN or PTEN-Nedd8 plasmid was transfected into the MCF-7 cells and immunoblotted with the indicated antibodies. **N** Coimmunoprecipitation of ITCH and PTEN after immunoprecipitation of the JUND protein in palbociclib sensitivity and resistance MCF-7 cells.

To date, no specific ubiquitin ligase for JUND has been reported. Through UbiBrowser 2.0 [42, 43], ITCH (also known as AIP4) was predicted to be the E3 of JUND with the highest affinity among the interactive protein candidates (Supplementary Fig. 2G). As expected, overexpression of ITCH reduced the levels of JUND protein and its phosphorylation, while *ITCH* knockdown showed opposing effects; however, this effect diminished following the onset of palbociclib resistance (Fig. 4F, G). Overexpression the wild-type of ITCH but not the ligase-deficient ITCH C830A mutant [44] promoted ubiquitination of JUND (Fig. 4H and Supplementary Fig. 2H). Similarly, knockdown of *ITCH* reduced JUND ubiquitination (Fig. 4I). Considering that neddylated PTEN in the nuclear intended to regulate the protein-protein interactions, we first used molecular docking with AlphaFold 2.0 to evaluate whether neddylation of PTEN affects the interaction between ITCH and JUND. The results indicated that neddylated PTEN could occupy the binding region between ITCH and JUND, whereas PTEN without Nedd8 could not (Fig. 4J). We then evaluated the interaction between ITCH and JUND and found that JUND was unable to bind to ITCH in sh*PTEN*+PTEN-Nedd8 MCF-7 cells with PTEN-Nedd8 overexpression (Fig. 4K-M). Furthermore, JUND failed to interact with ITCH in the palbociclib-resistant breast cancer cells compared to the palbociclib-sensitive cells (Fig. 4N). In addition, we noticed that PTEN neddylation increased the mRNA or protein levels of JUND, but not ITCH (Supplementary Fig. 2I). It is known that the proto-oncogene *JUN* is positively autoregulated by its product, JUN/AP-1 [45]. We proposed that PTEN neddylation enhanced JUND protein levels, which established positive feedback loop on the transcriptional level of *JUND* gene. Collectively, the results indicate that during drug resistance, elevated PTEN neddylation interferes with the interaction between JUND and its ubiquitin ligase, reducing JUND ubiquitination and increasing its protein stability.

### Identification of Echinacoside as a PTEN neddylation inhibitor

The above data indicate that PTEN neddylation is a key factor contributing to resistance against palbociclib treatment. Next, we aimed to explore the possibility of disrupting the interaction between PTEN and XIAP to reduce PTEN neddylation in tumor cells, and thus overcoming resistance to palbociclib. Consistent with previous study [30, 46], XIAP was co-immunoprecipitated with PTEN *in vivo* (Supplementary Fig. 3A). The truncations containing the BIR2 domain of XIAP and the C2 domain of PTEN interacted with each other (Fig. 5A and Supplementary Fig. 3B-D). Then, molecular docking and *in vitro* pull-down assay indicated that Y154 and N234 sites at XIAP, as well as S207 and R335 sites at PTEN were key binding sites on the interaction surface (Fig. 5B, C). On the basis of the binding determinants of the XIAP-PTEN interface, structure-based virtual screening was performed to find potential XIAP-PTEN PPI (protein and protein interaction) inhibitors (Fig. 5D). Ultimately, we selected top five compounds, guided by the top-ranking binding models and docking scores from the Enamine libraries containing protein-protein interacting compounds and the Bioactive Compound Library Plus at *Med Chem Express* LLC (Supplementary Fig. 3E). No PPI inhibitors affected the protein and mRNA levels of PTEN (Supplementary Fig. 3F, G). All the five compounds inhibited PTEN neddylation, with Echinacoside (ECH) and Z219118374 showing the most significant inhibition effect (Supplementary Fig. 3H). We noticed that Echinacoside (ECH), a natural phenylethanoid glycoside, had the highest molecular docking score and formed hydrogen bonds with the critical PTEN residues S207 and R335 that interacted with XIAP (Fig. 5E, F). ECH is one of the main phenylethanol glycosides isolated and purified from *cistanche tubulosa*, which has been used as traditional Chinese herbal medicine with anti-senility and antifatigue effects [47, 48]. Subsequently, we employed surface plasmon resonance and microscale thermophoresis assay to evaluate the interaction between ECH and PTEN. As illustrated in Fig. 5G and 5H, ECH showed a high-affinity binding with PTEN. In addition, TSA assay showed that the inflection temperature (Ti) of GST-PTEN increased after co-incubation with ECH (Supplementary Fig. 3I). Next, we monitored ECH binding with PTEN using the cellular thermal shift assay. The obvious shift in the melting curve indicated increased PTEN protein stability and high-affinity binding upon addition of ECH (Fig. 5I). Furthermore, strengthened XIAP-PTEN interaction was found under the drug resistance (Fig. 5J, K). However, ECH effectively disrupted the interaction between XIAP and PTEN in both palbociclib-sensitive and palbociclib-resistant cells *in vivo.* (Fig. 5J, K and Supplementary Fig. 3J). The pull-down assay revealed a clear dose-dependent decrease in the interaction between XIAP and the PTEN C2 domain following ECH treatment. (Fig. 5L). Moreover, ECH did not affect the dimerization of PTEN (Supplementary Fig. 3K, L).

**Fig. 5.**
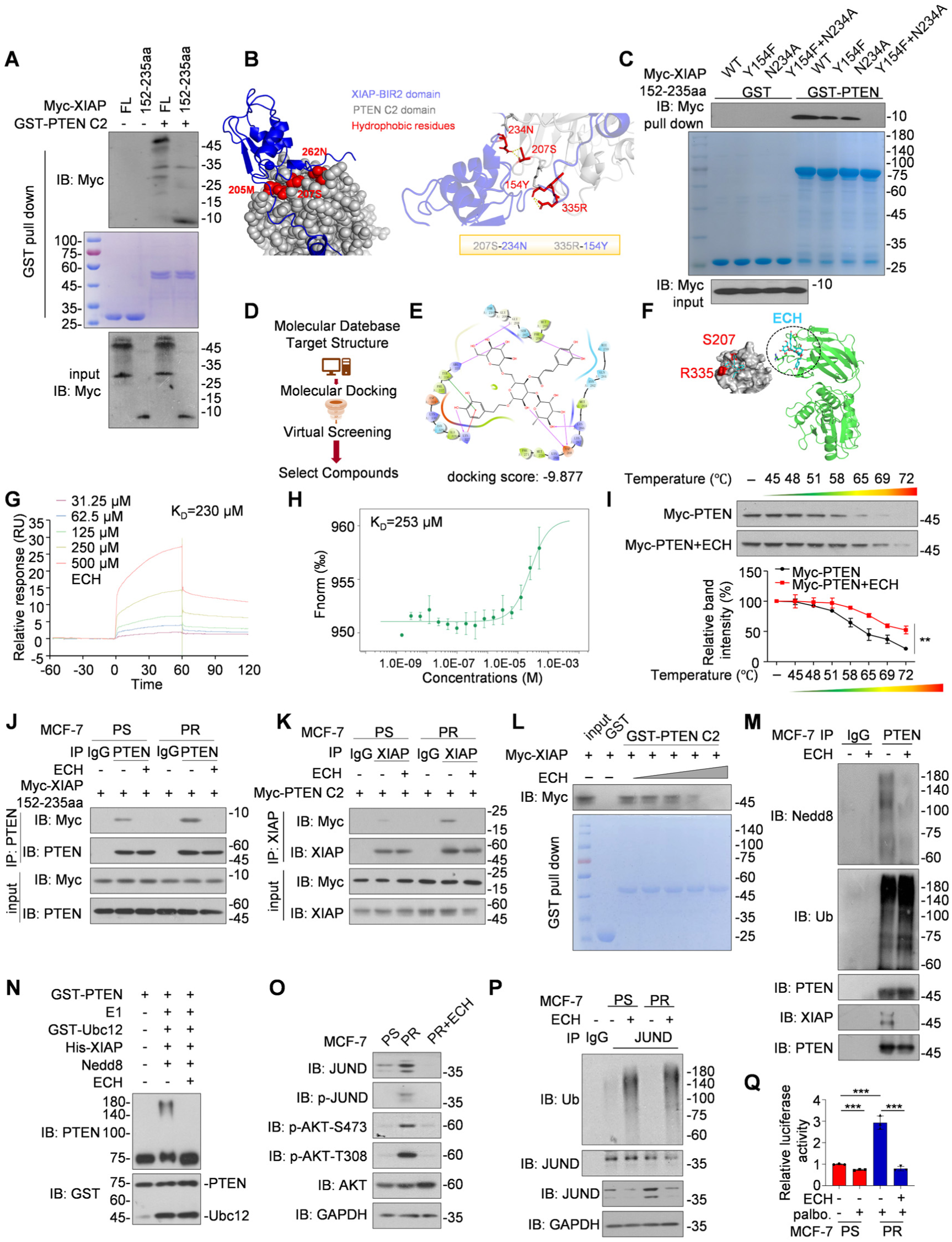
Echinacoside is identified as an inhibitor of PTEN neddylation. **A** Immunoblot analysis of GST pull-downs or WCL from HEK293T cells transfected with indicated constructs. **B** Molecular docking of PTEN C2 domain (gray) and XIAP BIR2 domain (blue). The key amino acids (red) at the PTEN and XIAP binding sites are shown. **C** Immunoblot analysis of GST pull-downs or WCL from HEK293T cells transfected with indicated constructs. **D** The workflow of virtual drug screening. **E** Structural formula and molecular docking score of Echinacoside with PTEN. **F** Molecular docking of Echinacoside (ECH) with PTEN was conducted to estimate the binding affinity. **G** Representative images of SPR binding assay of ECH to PTEN. ECH exhibited dose-dependent binding to GST-tagged PTEN that was immobilized onto a CM5 sensor chip surface. **H** All hits were titrated at various serially diluted concentrations (0.0153 μM to 500 μM) into a fixed concentration of GST-PTEN C2 domain (150 nM). **I** Dose and temperature-dependent CETSA experiments were conducted to verify the interaction between Echinacoside (ECH) and PTEN (n=3). The samples were divided into seven equal parts and heated at temperatures of 45 °C, 48 °C, 51 °C, 58 °C, 65 °C, 69 °C, and 72 °C. The abundance of PTEN protein was quantified using ImageJ. **J** Immunoprecipitates of PTEN were obtained from MCF-7-PS and MCF-7-PR cells transfected with Myc-XIAP (amino acids 152-235) and treated with Echinacoside (200 μM for 24 h). **K** Immunoprecipitates of XIAP were obtained from MCF-7-PS and MCF-7-PR cells transfected with Myc-PTEN C2 domain and treated with Echinacoside (200 μM for 24 h). **L** Immunoblot analysis of GST pull-downs or WCL from HEK293T cells transfected with indicated constructs. The concentration of ECH was used from 20 μM to 200 μM for 24 h. **M** PTEN neddylation was attenuated by ECH *in vivo*. The cells were treated with ECH at a concentration of 200 μM for 24 h. Immunoblot analysis was performed using anti-Nedd8, anti-Ub, anti-XIAP, and anti-PTEN antibodies on immunoprecipitates and whole cell lysates (WCL) from MCF-7 cells. **N** *In vitro* neddylation of PTEN. Purified His-XIAP and GST-PTEN proteins were incubated with Nedd8, Nedd8-E1/E2. Reactions were performed as described in the materials and methods section. Samples were analyzed by western blot with indicated antibodies. ECH (100 μg) was added and reacted for 1 h. **O** Immunoblot of WCL from MCF-7 PS and PR cells for indicated proteins. The cells were treated with 200 μM ECH for 24 h. **P** *In vivo* ubiquitination of PTEN. Immunoblot analysis of anti-JUND immunoprecipitate and WCL from MCF-7-PS cells and MCF-7-PR cells. **Q** AP-1 luciferase reporter plasmid was transfected in the palbociclib sensitivity and resistance MCF-7 cells. Reporter activity was assayed and represented as the means ± S.D. of three separate experiments. The cells were treated with ECH (200 μM) for 24 h.

To determine whether ECH could disrupt PTEN neddylation, we performed *in vivo* and *in vitro* neddylation assays. The results revealed that ECH reduced PTEN neddylation, but not ubiquitination (Fig. 5M). *In vitro* neddylation assay indicated that ECH directly interfered XIAP mediated PTEN neddylation (Fig. 5N). As expected, elevated JUND levels, JUND phosphorylation, and Akt phosphorylation, which increased after palbociclib resistance, were diminished following ECH treatment (Fig. 5O). By inhibiting PTEN neddylation, ECH reduced JUND mRNA levels, protein expression, and phosphorylation in a concentration-dependent manner, while exhibiting no effect on ITCH (Supplementary Fig. 3M, N). Additionally, there was a significant increase in JUND ubiquitination and decrease in AP-1 transcriptional activity following ECH treatment (Fig. 5P, Q). Taken together, we identified Echinacoside as a potent PTEN neddylation inhibitor that eliminates PTEN neddylation by inhibiting the interaction between XIAP and PTEN.

### ECH depends on PTEN neddylation to suppress tumorigenesis

Then, we intended to explore the role of ECH in tumorigenesis. ECH exhibited a significant tumor-suppressive effect, manifested in a dose-dependent manner, on cellular proliferation, colony formation, and cell migration across various tumor cell lines (Supplementary Fig. 4A-D). We measured drug toxicity of ECH in CD-1 mice and found no significant differences in body weight (Supplementary Fig. 5A, B), blood cell counts or composition (Supplementary Fig. 5C), or serum ALT levels (Supplementary Fig. 5D), indicating no obvious liver injury. Additionally, ECH treatment showed no signs of toxicity or organ damage based on weight and histopathology of vital organs (Supplementary Fig. 5E-G). Thus, we speculated that ECH might have substantial anti-tumor efficacy with minimal adverse effects.

To be note, deletion of *PTEN* significantly weakened the inhibitory effects of ECH on cell proliferation (Fig. 6A, B) and tumor invasion (Fig. 6C). Dose-response curves results showed that PTEN neddylation deficient cells were more sensitive to palbociclib than the sh*PTEN*+PTEN-WT cells (Fig. 6D). Moreover, palbociclib inhibited cell proliferation (Fig. 6E, F) and increased the cell death (Fig. 6G) in PTEN neddylation deficient cells. However, ECH was unable to inhibit tumor cell proliferation when PTEN neddylation was deficient (Fig. 6H, I). Additionally, in PTEN neddylation-deficient cells, ECH no longer reduced the protein levels of JUND and p-JUND (Fig. 6J), nor did it decrease AP-1 activity (Fig. 6K). Furthermore, CUT&Tag analysis performed on PTEN neddylation-deficient cells indicated an enrichment of the PI3K/Akt signaling pathway and the MAPK/AP-1 signaling pathway (Fig. 6L, M). The data demonstrated that in the absence of PTEN neddylation, JUND exhibited reduced enrichment at the promoters of downstream genes associated with inflammation and immune responses, including CSF2, IL-17, IL-6, MMP13, and IL-1β (Fig. 6N). Due to the significant changes in these genes, an immunoassay array was performed. As shown in Fig. 6O and 6P, the results demonstrated a reduction in multiple cytokines and chemokines, including G-CSF, GM-CSF, IFN-γ, IL-1RA, IL-1β, IL-8, CCL2/4, IL-17, and VEGF, following ECH treatment. However, no alterations were observed in the PTEN neddylation deficient cells subsequent to ECH exposure. Similar results of G-CSF, IL-1β, IL-17 were also approved by ELISA assay (Fig. 6Q). Therefore, the above evidence suggested that ECH inhibited oncogenic effects, at least to some extent, by interfering with PTEN neddylation.

**Fig. 6.**
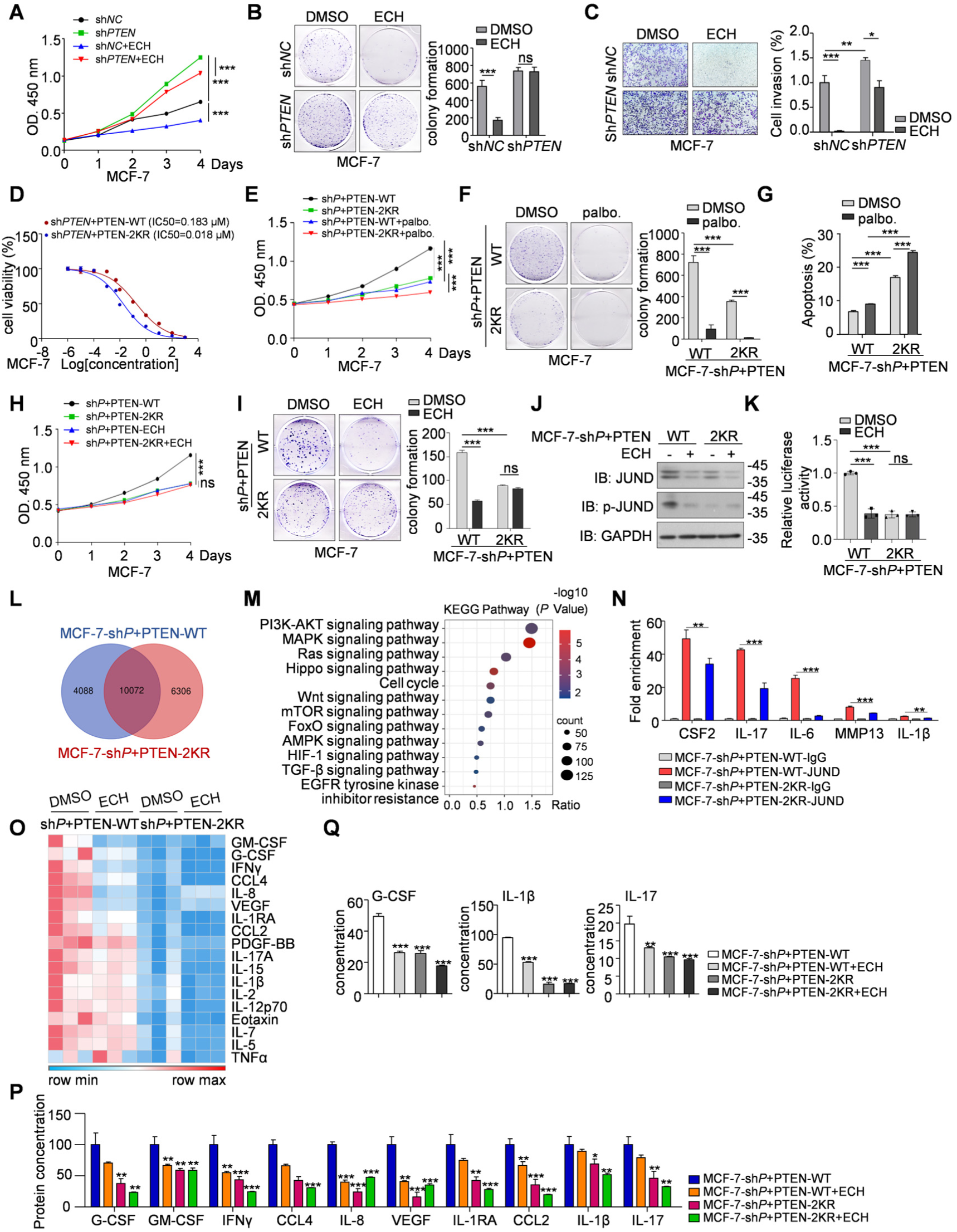
ECH is dependent on PTEN neddylation to suppress tumorigenesis. **A-C** CCK8 (**A**), colony formation (**B**) and cell invasion assay (**C**) were performed in the indicated MCF-7 cells treated with ECH. **D-G** CCK8 (**D, E**), colony formation (**F**) and apoptosis assay (**G**) were performed in the indicated MCF-7 cells treated with palbociclib. **H, I** CCK8 (**H**), and colony formation (**I**) were performed in the indicated MCF-7 cells treated with ECH. **J** Immunoblot of WCL from the sh*PTEN*+PTEN-WT/2KR MCF-7 cells treated by ECH (200 µM, 24 h) or not. **K** AP-1 luciferase reporter plasmid was transfected in the indicated cells. Reporter activity was assayed and represented as the means ± S.D. of three separate experiments. The cells were treated with ECH (200 μM) for 24 h. **L, M** Identification of transcription targets of the JUND in the sh*PTEN*-PTEN WT or neddylation deficient MCF-7 cells. Venn diagrams of overlapped promoters bound by JUND (**L**). A bubble chart of the enriched KEGG pathways comprising the 6306 target genes (**M**). **N** qChIP analysis of the indicated genes in indicated MCF-7 cells. Results are represented as fold change relative to control with GAPDH as a negative control. **O** Cytometric bead array was performed to detect the indicated cytokines. Heat map was performed to exhibit protein expression in the indicated cells. The cells were treated with ECH (200 μM) for 24 h. **P** The protein level of G-CSF, GM-CSF, IFNγ, CCL2, IL-8, VEGF, IL-1RA, IL-1β, IL-17 and CCL4 from the immunoassay array of the indicated MCF-7 cells are shown as bar chart. **Q** G-CSF, IL-1β and IL-17 protein levels were measured in the indicated MCF-7 cells by ELISA assay. *P* values were calculated by one-way ANOVA test (**A-C, E-I, K, N, P, Q**). Error bars, ± S.D. ns, not significant, \**P* < 0.05, \*\**P* < 0.01, \*\*\**P* < 0.001.

### Combination ECH enhances the sensitivity of palbociclib in breast cancer

We next aimed to explore whether ECH could overcome palbociclib resistance in breast cancer. As expected, the tumor weight and volume were both markedly declined in the palbociclib resistance MMTV-*PyMT* mice following the treatment of ECH (Fig. 7A-C). Similar results were also obtained from the primary breast cancer cells isolated from MMTV-*PyMT* mice (Supplementary Fig. 6A, B), and the tumor organoids established from them (Supplementary Fig. 6C-E). In immunoassay arrays, there was an obvious reduction of GM-CSF and G-CSF following ECH treatment (Fig. 6O, P). G-CSF is well known to function as a mobilizing agent for neutrophils, while GM-CSF influences the development and functionality of both neutrophils and macrophages [49–51]. Immunohistochemistry and flow cytometry showed a significant reduction in the population of active neutrophils labeled by Ly6G^+^, CD11b^+^, and CD62L^-^ in palbociclib-resistant MMTV-*PyMT* mice after ECH treatment (Fig. 7D, E). We confirmed a decrease in JUND, phosphorylation of Akt, PTEN neddylation, and XIAP levels, with no notable changes in neddylation of E1-UBA3, E2-Ubc12, and Nedd8 in palbociclib-resistant breast cancer tissues from MMTV-*PyMT* mice after ECH treatment (Supplementary Fig. 6F-I). The combination of ECH with palbociclib showed significantly decreased cell proliferation compared with monotherapy (Fig. 7F). In the ER-negative T98G cells, which was resistance to palbociclib, the co-administration of ECH increased sensitivity to palbociclib, thereby suppressing tumor cell proliferation (Supplementary Fig. 7A, B). We finally intended to confirm the efficacy of combination therapy using tumor organoid models derived from the human luminal breast cancer patient. Both ECH and palbociclib monotherapies demonstrated tumor growth suppression effects, but the combination treatment exhibited even better anti-tumor growth efficacy compared with the monotherapies (Fig. 7G, H). Immunofluorescence staining showed a significant reduction in PTEN neddylation following ECH treatment in the E-cadherin positive organoids (Fig. 7I). Taken together, these findings collectively illustrate that ECH serves as an effective sensitizing agent for palbociclib in breast cancer, with the combined therapy showing an enhanced anti-tumor activity.

**Fig. 7.**
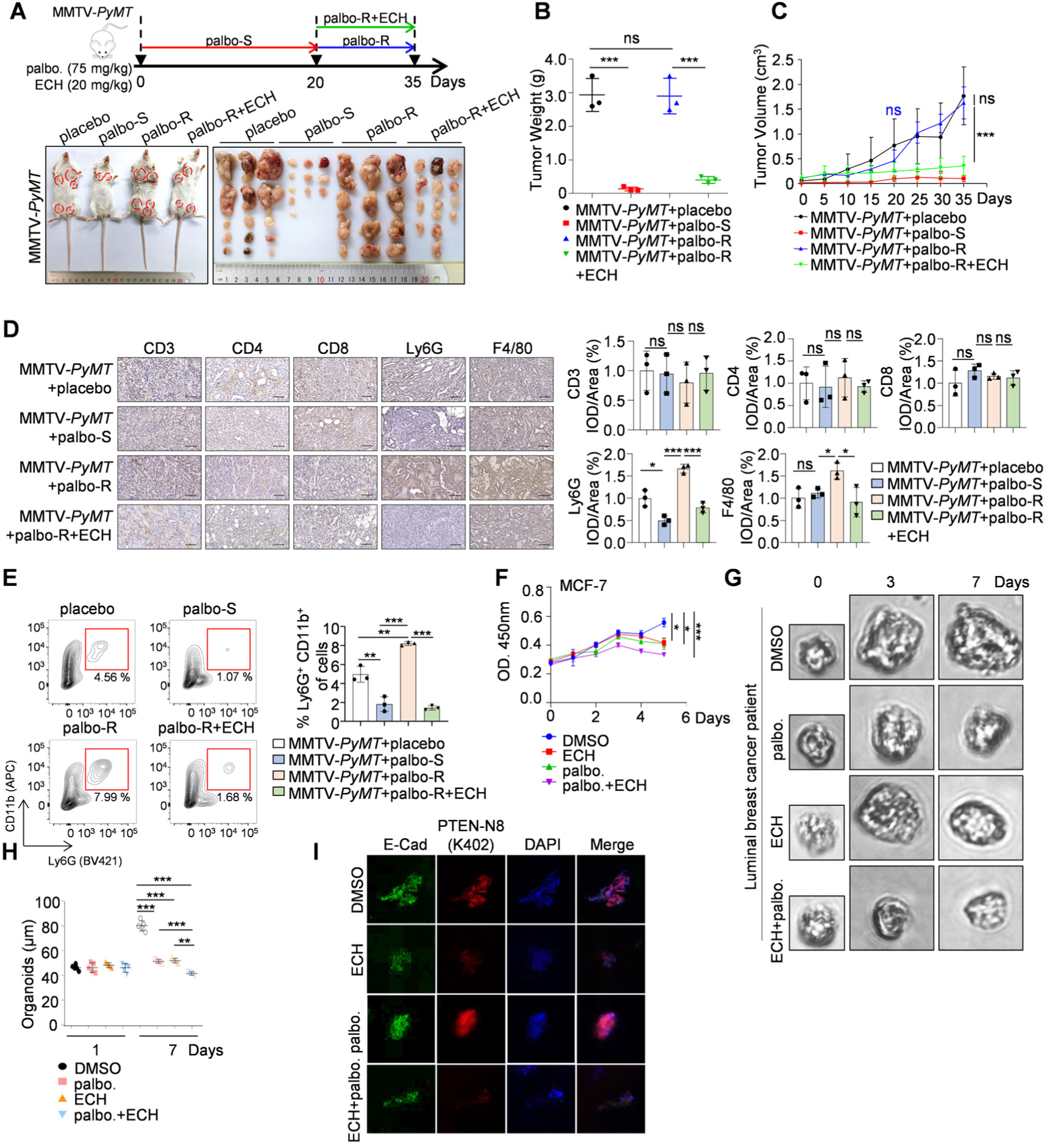
Combination ECH enhances the sensitivity of palbociclib in breast cancer. **A-C** The palbociclib-resistant mice (FVB/NJGpt background) model was established. The mice were treated with ECH (20 mg/kg) after palbociclib resistance for 15 days. The breast tumors were removed and photographed (n=3) (**A**). Tumors were isolated, then the weights (**B**) and their volume (**C**) were measured. Data are presented as means ± S.D. **D** Representative images from immunohistochemical staining of CD3, CD4, CD8, Ly6G and F4/80 in the indicated tumors. ImageJ was used to perform Semi-quantitative analysis. Scale bars, 50 µm. **E** Primary breast cancer cells from the indicated MMTV-*PyMT* mice were isolated and labeled with CD11b^+^ and Ly6G^+^ following flow cytometry sorting. **F** CCK8 assay was performed in MCF-7 cells treated with palbociclib (0.5 μM, 5 days) and ECH (200 μM, 5 days). **G, H** Representative images of organoids derived from a luminal breast cancer patient (**G**). Echinacoside (ECH) (200 μM) and palbociclib (0.5 μM) were administered either individually or in combination for 7 days. The diameters of the organoids are presented as a scatter plot (n=5) (**H**). **I** Immunofluorescent of E-Cadherin (green) and neddylated PTEN (K402) (red) in luminal breast cancer organoids. Data are shown as means ± S.D. and analyzed by a one-way ANOVA test (**B-F, H)**. ns, not significant, \**P* < 0.05, \*\**P* < 0.01, \*\*\**P* < 0.001.

## DISCUSSION

For luminal breast cancer patients, the standard treatment consists of CDK4/6 inhibitors combined with endocrine therapy. However, since resistance to CDK4/6 inhibitors is inevitable, there is considerable focus on developing therapeutic strategies to overcome this resistance. Here, we demonstrate that a post-translational modification of PTEN, namely neddylation, contributes to CDK4/6 inhibitor resistance in breast cancer. We firstly prepared antibodies that specifically recognize PTEN neddylation modification. Then, increased PTEN neddylation was observed in breast cancer patients resistant to palbociclib, as well as in both palbociclib-resistant breast cancer cells and spontaneous breast cancer MMTV-*PyMT* mice. Our previous data defined neddylation as a critical modification of PTEN, which inhibits PTEN tumor suppressor function and activates the PI3K/Akt signaling pathway, revealing a previously unidentified tumor-promoting role of neddylated PTEN [30]. Palbociclib resistance upregulated PTEN neddylation *via* the E3 ligase XIAP, consistent with previous studies suggesting that compensatory CDK2 activation upon resistance increases XIAP mRNA levels [37, 38]. This dynamic resulted in enhanced binding to PTEN, thereby contributing to the augmented PTEN neddylation and hyperactivated PI3K/Akt signaling pathway observed following palbociclib resistance.

*PIK3CA* mutations occur in approximately 40 % of luminal breast cancers, and activation of the PI3K/Akt signaling pathway is prominent as cancers become resistant to CDK4/6 inhibitors [15]. While PI3K inhibitors hold promise in mitigating the onset of CDK4/6 inhibitor resistance, their efficacy in rescuing cells from established resistance falls short, as they are unable to reverse the acquired insensitivity [52–54]. Apart from the upregulation of PI3K/Akt signaling pathway, our study showed that the increase in PTEN neddylation after palbociclib resistance led to the over-activation of the MAPK/AP-1 signaling pathway. Mechanistically, we identified that neddylated PTEN interacted with JUND. JUND is a component protein of the activating protein-1 (AP-1) transcription factor and is considered as a classical proto-oncogene. Dysfunctional AP-1 transcriptional activity is associated with tumor growth, invasion, migration and drug resistance [39, 55, 56]. So far, it is known that CDK4/6 inhibition reprograms the breast cancer enhancer landscape by stimulating AP-1 transcriptional activity [57]. However, the role of AP-1 in regulating resistance to CDK4/6 inhibitors remains unclear. Here, we reported that PTEN neddylation led to an obviously enhancement in JUND protein stability in palbociclib resistance breast cancer cells and mouse models. The expression of JUND in palbociclib-resistant patients showed a positive correlation with the PTEN neddylation modification level and a negative correlation with PTEN expression. Up to day, no ubiquitin ligase has been clearly reported to catalyze the ubiquitination mediated degradation of JUND. We identified that ITCH as a polyubiquitination E3 ligase for JUND. However, under the palbociclib-resistant conditions, ITCH no longer promoted JUND ubiquitination and reduced its protein stability. Further structural docking and immunoprecipitation assays showed that neddylated PTEN interfered with the interaction between JUND and ITCH, reducing the ubiquitination of JUND catalyzed by ITCH, thereby preventing the degradation of JUND. Therefore, the increased modification of PTEN neddylation after palbociclib resistance enhanced the protein stability and phosphorylation level of JUND, thereby activating the AP-1 transcription factor activity. Thus, further data showed that the expression levels of JUND downstream target genes such as CSF2, IL-1β, IL-6, IL-17, and MMP13 were increased. These results propose a novel role for PTEN neddylation in upregulating AP-1 transcription factor activity and activating the MAPK/AP-1 signaling pathway during resistance to palbociclib treatment.

Neddylation is indispensable in various cellular processes and homeostasis, but excessive activation of this modification has been observed in numerous tumors. Small molecules targeting neddylation activating enzymes (E1), such as MLN4924 (trade name pevonedistat) and TAS4464, have shown promise in clinical trials for cancer treatment [28, 58]. DI-404, DI-591, NAcM-HIT, NAcM-OPT, and WS-383 are competitive neddylation conjugating enzymes (E2) inhibitors that target UBE2M-DCN1 interaction [59–61]. HA-9104 is a targeted inhibitor of another E2-UBE2F [62]. Given the limited number of neddylation E1 and E2 enzymes, inhibiting neddylation broadly may pose inevitable cytotoxicity and drug safety concerns. Therefore, for tumor treatment, more specific and targeted drugs are needed to address neddylation modifications. No small molecule drugs specifically targeting neddylation substrates have been reported, but we believe that would offer greater specificity, reduced drug side effects, and potentially more potent anti-tumor effects. XIAP has been identified as a specific ligase for PTEN neddylation both *in vivo* and *in vitro* [30]. Next, we employed a specific small molecule inhibitor, Echinacoside (ECH), to interfere with the dynamic interplay between XIAP and PTEN, ultimately suppressing the neddylation of PTEN both *in vivo* and *in vitro*. ECH, a key efficacious constituent found in *Echinacea*, is also abundantly present in various other natural plant sources, including the roots of *Scrophulariae*, *Rehmanniae*, and *Cistanches Herba*, among others [45, 46]. It has been reported that ECH inhibits tumor metastasis *via* suppressing the PI3K/AKT signaling pathway, but the mechanism remains unclear [63–65]. Our research demonstrates that ECH exhibits excellent anti-tumor efficacy through its effect on PTEN neddylation. For the first time, our study demonstrates that ECH is a selective inhibitor targeting neddylation substrates.

Although the resistance mechanisms associated with CDK4/6 inhibitors in breast cancer have been documented, but not fully elucidated. For instance, the concurrent use of PI3K inhibitors like alpelisib or buparlisib with CDK4/6 inhibitors is discouraged due to safety concerns [14, 15]. Since PTEN is a key suppressor in the PI3K/Akt pathway, restoring its tumor-suppressive function could more effectively inhibit this pathway [23]. More importantly, our data reported that ECH obviously elevates the sensitivity of palbociclib, and ECH in combination with palbociclib would more effectively reduce the tumor growth. In the tumor microenvironment, the role of neutrophils is complex and multifaceted, highly correlated with the development of drug resistance [66, 67]. Our data showed that the number of activated neutrophils increases after palbociclib resistance, accompanied by an increase in various pro-inflammatory factors and growth factors, such as G-CSF, GM-CSF, IL-1β, VEGF. Treatment with ECH was observed to decrease activated neutrophils following palbociclib resistance. Future study needs to explore the role of activated neutrophils in ROS production and the immunosuppressive microenvironment, and discuss the function of other different terms of neutrophils. Ultimately, our findings unlock a promising therapeutic strategy by focusing on neddylation to effectively restore PTEN tumor suppressor activity, thereby enhancing the efficacy of palbociclib. It highlights a deeper understanding of CDK4/6i resistance mechanisms and the development of advanced combination therapies in optimizing breast cancer treatment outcomes.

## METHODS AND MATERIALS

### Patients and mice

The clinical samples were approved by the department of breast and thyroid surgery at the Second People’s Hospital of Shenzhen and Hunan Cancer Hospital. Informed consent was obtained from all subjects or their relatives. The MMTV-*PyMT* mice (*PyMT*/WT mice, female, background C57BL/6J (RRID: IMSR_JAX: 000664)) were purchased from Cyagen Biosciences. The MMTV-*PyMT* mice (*PyMT*/WT mice, female, background FVB/NJGpt (RRID: IMSR_GPT: T004993)) were kindly gifted by the Second People’s Hospital of Shenzhen. CD-1 (ICR, RRID: IMSR_CRL: 022) mice were purchased from Beijing Vital River Laboratory Animal Technology Co., Ltd. Animals were assigned to experimental groups using simple randomization.

### Cell culture and reagents

All mammalian cells were cultured following ATCC recommended protocols and in medium supplemented with 10 % fetal bovine serum (FBS, VivaCell BIOSCIENCES C04001-500) and 1 % Penicillin-Streptomycin (Gene-Protein Link P08X14). MCF-7 (RRID: CVCL_0031), HEK293T (RRID: CVCL_0063), HCT116 (RRID: CVCL_0291), A549 (RRID: CVCL_0023), MDA-MB-231 (RRID: CVCL_0062), MCF-7 sh*PTEN*, MCF-7 sh*PTEN*+PTEN-Nedd8, MCF-7 sh*PTEN*+NLS-PTEN, MCF-7 sh*PTEN*+PTEN-WT and MCF-7 sh*PTEN*+PTEN-2KR cells were cultured in Dulbecco’s modified Eagle’s medium (DMEM) with high glucose (Thermo Fisher Scientific). T-47D (RRID: CVCL_0553) and T98G (RRID: CVCL_0556) cells were cultured in RPMI-1640 medium (Thermo Fisher Scientific C11875500BT). MCF-7 sublines resistant to doxorubicin (MCF-7/ADR) were purchased from Sunncell Biosciences and were cultured in RPMI-1640 medium supplemented with 500 ng/ml Doxorubicin hydrochloride. MCF-7 sublines resistant to cisplatin (MCF-7/DDP) were purchased from Shanghai Mei Xuan Biological Technology Co., Ltd and were cultured in RPMI-1640 medium supplemented with 500 ng/ml Cisplatin. MCF-7 sublines resistant to paclitaxel (MCF-7/PTX) were purchased from BLUEFBIO (Shanghai, China) and were cultured in Dulbecco’s modified Eagle’s medium (DMEM) with high glucose supplemented with 200 ng/ml Paclitaxel. All cell lines were authenticated by STR profiling.

Palbociclib (HY-50767), Pevonedistat (HY-70062), Echinacoside (HY-N0020), Maltotetraose (HY-N2464), MG132 (HY-13259), Cisplatin (HY-17394), Paclitaxel (HY-B0015), Doxorubicin hydrochloride (HY-15142), Z21911837, Z45457765 and Z1549417413 were purchased from *Med Chem Express*. *ITCH* and negative control small interfering RNA (siRNA) were acquired from GenePharma (Suzhou GenePharma Co.,Ltd). The sequences of siRNAs are shown in Supplementary Table 4.

### Establishment of palbociclib-resistant cells

To generate the palbociclib-resistant cells, the cells were cultured in media supplemented with increasing concentrations of palbociclib for over 6 months. The concentration of palbociclib were began at 100 nmol/L and were maintained at a final dose of 6 μmol/L, and the palbociclib-resistant cells were named as MCF-7-PR cells (MCF-7 palbociclib-resistant cells). For all experiments, the resistant cells were refreshed with palbociclib-free media for 48 h before treatment.

### Antibodies

The following antibodies were used: Phospho-Rb (Ser807/811) (Cat# 8516, RRID: AB_11178658), Rb (Cat# 9309, RRID: AB_823629), Cyclin D1 (Cat# 2922, RRID: AB_2228523), PTEN (Cat# 9188, RRID: AB_2253290), Ubiquitin (Cat# 20326, RRID: AB_3064918), Akt (Cat# 9272, RRID: AB_329827), Phospho-Akt (Ser473) (Cat# 9271, RRID: AB_329825), Phospho-Akt (Thr308) (Cat# 13038, RRID: AB_2629447), Phospho-JUND (Ser100) (Cat# 9164, RRID: AB_330892), CUL1 (Cat# 4995, RRID: AB_2261133), XIAP (Cat# 2042, RRID: AB_2214870), Nedd8 (Cat# 2754, RRID: AB_659972), c-JUN (Cat# 9165, RRID: AB_2130165), JUNB (Cat# 3753, RRID: AB_2130002), c-FOS (Cat# 2250, RRID: AB_2247211) and FOSB (Cat# 2251, RRID: AB_2106903) antibodies were purchased from Cell Signaling Technology. Ly6G (Cat# sc-53515, RRID: AB_783639), CD3 (Cat# sc-18843, RRID: AB_627010), CD4 (Cat# sc-53042, RRID: AB_629062), CD8 (Cat# sc-53063, RRID: AB_629231), CD11b (Cat# sc-1186, RRID: AB_626884), NEDP1 (Cat# sc-271498, RRID: AB_10649498) and Cyclin E (Cat# sc-377100, RRID: AB_2923122) antibodies were purchased from Santa Cruz Biotechnology. F4/80 (Cat# 28463-1-AP, RRID: AB_2881149), GAPDH (Cat# 60004-1-Ig, RRID: AB_2107436), UBA3 (Cat# 68067-1-Ig, RRID: AB_2918808), Ubc12 (Cat# CL488-67482, RRID: AB_3084351), ITCH (Cat#20920-1-AP, RRID: AB_2878765) and GST (Cat# HRP-66001, RRID: AB_2883833) antibodies were purchased from Proteintech. Flag (Cat# M185-3L, RRID: AB_11123930), Myc (Cat# M047-3, RRID: AB_591112) and HA (Cat# M132-3, RRID: AB_10207271) antibodies were purchased from MBL. JUND (Cat# A5433, RRID: AB_2863501) and FOSL1 (Cat# A5372, RRID: AB_2766182) antibodies were purchased from ABclonal. Goat anti-Rabbit IgG (H + L) Secondary Antibody (Cat# 65-6120, RRID: AB_2533967) and Goat anti-Mouse IgG (H + L) Secondary Antibody (Cat# 31430, RRID: AB_228307) were purchased from Thermo Fisher Scientific. PTEN-Nedd8 (K402) antibody was purchased from PTM BioLab (Hang Zhou) Co., Inc.

### Establishment of palbociclib-resistant mice

The palbociclib-sensitive MMTV-*PyMT* mice were obtained by continuing palbociclib (75 mg/kg or 100 mg/kg) gavage for 20 days. Palbociclib-resistant MMTV-*PyMT* mice were obtained by continuing palbociclib (75 mg/kg or 100 mg/kg) gavage for 35 or 50 days.

### Immunoprecipitation and immunoblotting

For immunoprecipitation assays, cells were lysed in EBC lysis buffer (0.5 % NP-40, 50 mM Tris, pH 7.6, 120 mM NaCl, 1 mM EDTA, 1 mM Na_3_VO_4_, 50 mM NaF and 1 mM β-mercaptoethanol) supplemented with protease inhibitor cocktail (Roche, 11836170001). Immunoprecipitations were performed using the indicated primary antibody and Dynabeads™ protein G beads (10004D) at 4 °C. The immunoprecipitants were washed at least three times in EBC lysis buffer before being resolved by SDS-PAGE and immunoblotted with indicated antibodies.

### *In vivo* modification assays

Cells were solubilized in modified lysis buffer (50 mM Tris, pH 7.4, 150 mM NaCl, 10 % glycerol, 1 mM EDTA, 1 mM EGTA, 1 % SDS, 1 mM Na_3_VO_4_, 1 mM DTT and 10 mM NaF) supplemented with a protease inhibitor cocktail. The cell lysate was incubated at 60 °C for 10 min. The lysate was then diluted 10 times with modified lysis buffer without SDS and incubated with the indicated antibody for 3 h at 4 °C before adding Dynabeads™ protein G beads (10004D). Then the lysate was rotated gently for 8 h at 4 °C. The immunoprecipitants were washed at least three times in wash buffer (50 mM Tris, pH 7.4, 150 mM NaCl, 10 % glycerol, 1 mM EDTA, 1 mM EGTA, 0.1 % SDS, 1 mM DTT and 10 mM NaF). Proteins were analyzed by western blotting with indicated antibodies.

### *In vitro* modification assays

*In vitro* neddylation was performed using recombinant purified enzymes. GST-PTEN and GST-Ubc12 were expressed in Escherichia coli BL21 (DE3). 0.5 μg of GST-PTEN and 0.5 μg of GST-Ubc12 were incubated with 2 μg of Nedd8, 10 ng of E1 (APPBP1 UBA3) and 0.5 μg of E3 (His-XIAP) in a total reaction volume of 30 μL Nedd8 conjugation Rxn Buffer Kit (Boston Biochem, SK-20). Nedd8 (UL-812), Nedd8 E1 (APPBP1/UBA3) (E-313), His-XIAP were purchased from Boston Biochem. Samples were incubated at 30 for 1 h, and reactions were terminated with 2 × SDS-PAGE loading buffer (20 mM Tris-HCl, pH 6.8, 100 mM DTT, 2 % SDS, 20 % glycerol and 0.016 % Bromophenol blue) before western blotting. If needed, the neddylation reaction sample was first purified with GST-tag Purification Resin.

### Dual Luciferase assay

MCF-7 cells were transfected using Lipofectamine 3000 (Invitrogen) according to manufacturer’s protocol. After 48 hours, cells were lysed in 100 μL of a passive lysis buffer (Promega). Luciferase activity was measured with the Dual Luciferase Assay System (Promega) according to the manufacturer’s protocol.

### GST pull down

The PTEN or PTEN C2 domain sequence was inserted into the pGEX-4T-3 vector (Amersham). To detect the direct binding, bacteria-expressed GST-tagged proteins were immobilized on BeyoGold™ GST-tag Purification Resin (P2250) and then incubated with Myc-tagged proteins for 8 h at 4 under rotation. Beads were washed with GST-binding buffer (100 mM NaCl, 50 mM NaF, 2 mM EDTA, 1 % NP-40 and protease inhibitor mixture) three times and proteins were eluted, followed by immunoblotting.

### Molecular docking and virtual screening

The molecular docking of PTEN (PDB ID5R) and XIAP (PDB 1C9Q) was performed using the reliable online docking server ZDOCK (http: //zdock.umassmed.edu/). The interaction between PTEN and XIAP was visualized using PYMOL version 1.7.0.3. All ten docking complexes were used for our analysis. The cryo-EM structure of human PTEN (PDB ID5R) was used to perform structure-based virtual screening via a workflow application of Glide in Maestro (Schrödinger Maestro 11.4). The Protein-Protein Interaction Library and the MCE Bioactive Compound Library Plus were selected as screening databases. The LigPrep module of Maestro was used to generate the three-dimensional (3D) conformations of the compounds. At the very beginning, the compounds were screened using the high-throughput virtual screening module and the top-ranked 20 % candidates were selected on the basis of the Glide G-score. Then these candidates were redocked using the standard precision and extra precision mode in turn. Finally, 200 compounds with shape rationality and structural diversity were picked for assay.

### Surface plasmon resonance binding assays (SPR)

Surface plasmon resonance experiments were performed on a BIACORE 3000 biosensor system (GE Healthcare) according to the manufacturer’s instructions at 25 °C. PTEN protein was immobilized onto a CM5 chip (GE Healthcare), which was activated using a 1: 1 mixture of 1-ethyl-3-(3-dimethyllaminopropyl) carbodiimide (EDC) and N-hydroxysuccinimide (NHS) at a flow rate of 10 μL/min. ECH was diluted in PBS, and then injected over the PTEN surface at a rate of 30 μL/min flow rate. The duration of protein binding time was set to 180 s, after which the running buffer was injected at the same rate for 300 s. The binding kinetics were processed and calculated by BIAevaluation software.

### Microscale thermophoresis assays (MST)

Microscale thermophoresis assays detected interactions between PTEN and ECH. The concentration of PTEN protein was held constantly at 150 nM, whereas the concentrations of ECH were gradient-diluted (0.0153 μM-500 μM). After brief incubation, the samples were loaded into MST-standard glass capillaries. The measurements were performed on a MST machine (NanoTemper Technologies, Munich, Germany).

### Cellular thermal shift assay (CETSA)

HEK293T cells (1 × 10^7^) which transfected Myc-PTEN were resuspended in PBS containing 1 % protease inhibitors. The cells were then lysed by liquid nitrogen through three cycles of freezing and thawing. The resulting cell lysates were centrifuged at 12 000 g for 15 min at 4 °C to collect the supernatants. Subsequently, the samples were divided into 7 equal parts and heated at temperatures of 45 °C, 48 °C, 51 °C, 58 °C, 65 °C, 69 °C, and 72 °C for 3 min each. After heating, the samples were centrifuged at 12 000 g for 15 min at 4 °C, and the resulting supernatants were collected and boiled with loading buffer. The levels of the target proteins were determined using western blotting.

### Fluorescence microscopy

After fixation with 4 % paraformaldehyde and permeabilization in 0.2 % Triton X-100 (PBS), cells were incubated with the indicated antibodies for 12 h at 4 °C, followed by incubation with goat anti rabbit IgG H&L Alexa Fluor 488 or 568 antibody for 1 h at 37 °C. The nuclei were stained with DAPI, and images were visualized with a Leica confocal microscopy.

### Tumor organoid culture

The breast cancer tissues were minced and digested with 5 mL of TrypLE (Gibco 12605-010) digestion solution at 37 °C for 1 hour. The digestion was stopped by adding 5 mL of culture medium, followed by centrifugation at 300 g for 5 minutes. The cell pellet was washed with 10 mL of PBS and then filtered through a 70 µm cell strainer. The filtered suspension was centrifuged again at 300 g for 5 minutes. Pre-chilled pipette tips were used to resuspend the bottom cells in 50 µL of pre-chilled organoid complete medium and keep them on ice. Then, 250 µL of Matrigel was added to the cell suspension. A pre-warmed 96-well plate was removed, and 30 µL of each drop of the cell suspension was aspirated into the well of the plate using a pre-chilled pipette tip. The plate was quickly inverted and placed in a 37 °C, 5 % CO incubator to allow the matrix gel to solidify for 15 minutes. After solidification, an appropriate volume of organoid complete medium was added to each well.

### CUT&Tag sequencing

CUT&Tag assay was performed using NovoNGS CUT&Tag 3.0 High-Sensitivity Kit (for Illumina, Novoprotein scientific Inc., Cat#N259-YH01-01A). Briefly, 1 × 10^5^ cells were harvested freshly. Then 37 % formaldehyde was gently added to cells and these samples were incubated at room temperature for 2 min. Cross-linking was stopped by addition of 1.25 M glycine and then the samples were washed with the wash buffer. The cells were enriched by ConA Beads and resuspended by 50 µL primary antibody buffer of anti-JUND antibody, and incubated overnight at 4 °C. Then discarded the primary antibody buffer and added 100 µL anti-rabbit IgG antibody buffer for 1 h at a dilution of 1: 100. Then the beads were washed for 3 times using antibody buffer and then incubated with protein A/G-Tn5 transposome for 1 h and washed for 3 times by ChiTaq buffer. Cells were resuspended in 50 µL Tagmentation buffer and incubated at 37 °C for 1 h. The incubation was stopped by adding 10 µL 10 % SDS at 55 °C for 10 min. The DNA fragments were extracted by Tagment DNA extract beads and amplified using 5 × AmpliMix. Then the DNA was re-extracted by DNA clean beads for sequencing.

### Chromatin immunoprecipitation (ChIP) and qChIP

ChIP assays were performed using the EZ-Magna ChIP A/G Chro matin Immunoprecipitation Kit (17-10086, Merck Millipore) according to the manufacturer’s recommendations. In brief, the harvested MCF-7 cells were cross-linked with 1 % formaldehyde at 37 °C for 20 min, lysed, and sonicated to generate 200 to 1000 bp DNA fragments. Antibodies against JUND and rabbit IgG were used for immunoprecipitation, and the precipitated DNA fragments were subjected to qPCR amplification. The primers used were listed in Supplementary Table 3.

### Flow cytometry analysis and FACS sorting

For detection of apoptosis, cells were co-stained with Annexin-V-phycoerythrin and amino actinomycin D (7-AAD) according to the manufacturer’s instructions. Stained cells were sorted with fluorescence-activated cell sorting (FACS). The isolated cell suspension was washed with PBS, then stained with specific mAbs at indicated concentration in PBS solution containing 1 % FBS. Antibody staining was performed at 4 °C for 30 min and the stained cells were analyzed on an LSRFortessa cell analyzer (BD Biosciences). FACS sorting routinely yielded cell purity levels of over 90 %. Data were analyzed using FlowJo software (RRID: SCR_008520).

### Cohort and immunohistochemistry

The breast tissue microarrays were purchased from Shanghai Biochip Company (HBreD090Bc03) and Shanghai Zhuoli Biotech Company (ZL-Brcsur1801). ZL-Brcsur1801 was immunostained with PTEN and Nedd8-PTEN (K402) antibodies (Fig. 1A-C). Detailed pathological information was in Supplementary Table 1. HBreD090Bc03 was immunostained with Nedd8-PTEN (K402) and JUND antibodies (Supplementary Fig. 2C-F), which are available in Supplementary Table 2. In brief, tissues were fixed with 4 % PFA and embedded in paraffin. The samples were then attached to slides, washed and incubated with 3 % H_2_O_2_ for 10 min. Afterwards, these sections were blocked and then incubated with a primary antibody at 4 °C overnight. After washing with PBS, the sections were incubated with secondary antibody for 1 h at room temperature. Finally, the sections were washed and incubated with DAB substrate for visualization. After washing and dehydration, the sections were sealed with coverslips. All staining was assessed by a quantitative imaging method; the percentage of immunostaining and the staining intensity were recorded. An H-score was calculated using the following formula: H-score = Σ (PI × I) = (percentage of cells of weak intensity × 1) + (percentage of cells of moderate intensity × 2) + (percentage of cells of strong intensity × 3). PI indicates the percentage of positive cells versus all cells.

### Multiplex immunohistochemical

Pheno 7-plex IHC kit (Pheno, PVB2002) was used according to manufacturer’s protocol. Slides were deparaffinised in xylene and rehydrated with ethanol and double distilled water. Antigen was retrieved by citrate buffer and heated to boil in microwave for approximately 20 min, sections were then blocked with bovine serum albumin at room temperature for 10 min. Primary antibodies were incubated at 37 °C for 30 min, secondary antibody conjugated with horseradish peroxidase was incubated at room temperature for 10 min. Tyramide signal amplification was performed with 1: 100 in antibody dilution buffer diluted fluorophores and incubated at room temperature for 10 min. Nuclei were stained with DAPI after primary antibodies were stained.

### ELISA assay

The quantitation of IL-1β, IL-17A and G-CSF levels in MCF-7 cells were performed using ELISA assay kit (Solarbio, SEKM-0002, SEKH-0026, and SEKH-0058) according to the manufacturer’s instructions.

### Immunoassay arrays

Cytokine concentrations of supernatants of indicated cells were measured using the CBA (Cytometric Bead Array, RayBiotech FAH-STRM-1) following the manufacturer’ s instruction. Read samples on a flow cytometer and analyze using FCAP Array software (BD Biosciences).

### Statistical analysis

All data were representative of no fewer than three independent experiments. All data were presented as means ± S.D. Data were analyzed using SPSS 19.0 (IBM Corp., Armonk, NY, USA, RRID: SCR_002865) and GraphPad Prism 8 (GraphPad Software, Inc., La Jolla, CA, RRID: SCR_002798). The differences between groups were assessed by Student’s *t*-test. The correlations were analyzed using the pearson correlation test. A value of *P* < 0.05 was considered to be statistically significant.

## DATA AVAILABILITY

RNA-seq data have been deposited at SRA (PRJNA1143082 and PRJNA1144658) and are publicly available as of the date of publication. All data are available in the main text or the supplementary materials.

## AUTHOR CONTRIBUTIONS

Conceptualization: PX; Methodology: FL, WXL, YXS, SHL, XKJ; Formal Analysis: FL; Investigation: PX, FL; Resources: SYS, YWT, ZZ; Writing-Original Draft: PX; Funding Acquisition: PX.

## ETHICS APPROVAL

All animals were handled in strict accordance to the “Guide for the Care and Use of Laboratory Animals” and the “Principles for the Utilization and Care of Vertebrate Animals”, and all animal work was approved by Capital Medical University Ethics Committee. The clinical samples were approved by the department of breast and thyroid surgery at the Second People’s Hospital of Shenzhen and Hunan Cancer Hospital. Informed consent was obtained from all subjects or their relatives.

## ACKNOWLEDGEMENTS

Thanks to Core Facility Center, Capital Medical University for providing platform support for this research.

