## Supplementary material for "PTEN neddylation aggravates CDK4/6 inhibitor resistance in breast cancer": Supplementary information-clean version.pdf

Fan Liu *et al.*

**This PDF file includes:**

Supplementary Figs. 1-7

Supplementary methods

**Other Supplementary Materials for this manuscript include the following:**

Supplementary Table 1. List of pathological information of ZL-Brcsur1801.

Supplementary Table 2. List of pathological information of HBreD090Bc03.

Supplementary Table 3. Primers.

Supplementary Table 4. Sequences for RNA interference.

**A**

| Sample number | Gender | Age | Receptor status | Ki-67 | Grade |
|---|---|---|---|---|---|
| **Preinvasive breast carcinoma** | | | | | |
| 1 | Female | 65 | ER+/PR+/HER2- | + | Ⅲ |
| 2 | Female | 65 | ER+/PR+/HER2- | + | Ⅱ |
| 3 | Female | 53 | ER+/PR+/HER2- | + | Ⅲ |
| 4 | Female | 48 | ER+/PR+/HER2- | + | Ⅲ |
| 5 | Female | 43 | ER+/PR+/HER2- | + | Ⅱ |
| 6 | Female | 38 | ER+/PR+/HER2- | + | Ⅲ |
| 7 | Female | 57 | ER+/PR+/HER2- | + | Ⅱ |
| **Recurrent breast carcinoma** | | | | | |
| 8 | Female | 46 | ER+/PR+/HER2- | + | ⅢA |
| 9 | Female | 64 | ER+/PR+/HER2- | + | Ⅳ |
| 10 | Female | 36 | ER+/PR+/HER2- | + | Ⅱ-Ⅲ |
| 11 | Female | 41 | ER+/PR+/HER2- | + | Ⅱ |

**B**

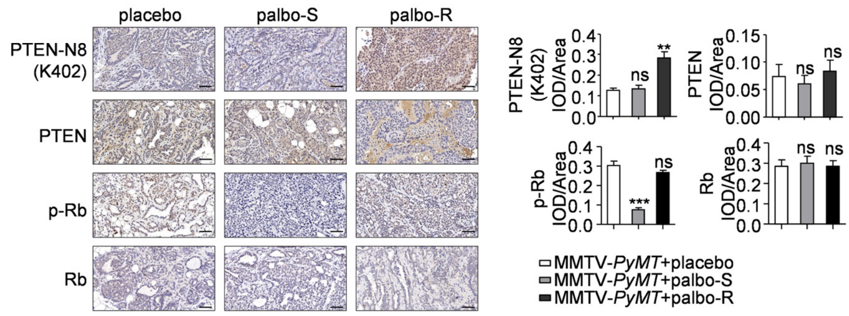


**C**

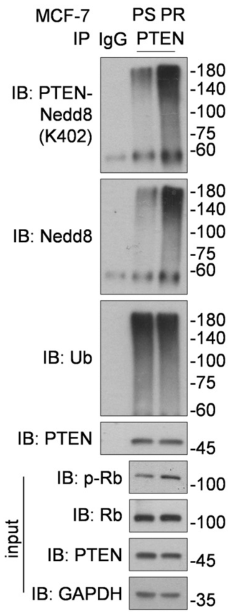


**Supplementary Fig. 1 PTEN neddylation modification is upregulated in the CDK4/6 inhibitor-resistant breast cancer. A** Pathological information of patients with preinvasive breast cancer, and recurrent breast cancer after palbociclib treatment. **B** Representative images from immunohistochemical staining of PTEN, PTEN neddylation on K402, Rb, p-Rb in control group, palbo-sensitive (palbo-S) and palbo-resistant (palbo-R) tumors. ImageJ was used to perform Semi-quantitative analysis. Scale bar, 50 µm. Data were analyzed using the one-way ANOVA test. **C** *In*

*vivo* neddylation and ubiquitination modification assay. Immunoprecipitation of PTEN in the MCF-7-PS and MCF-7-PR cells and analysis using immunoblotting with the indicated antibodies.

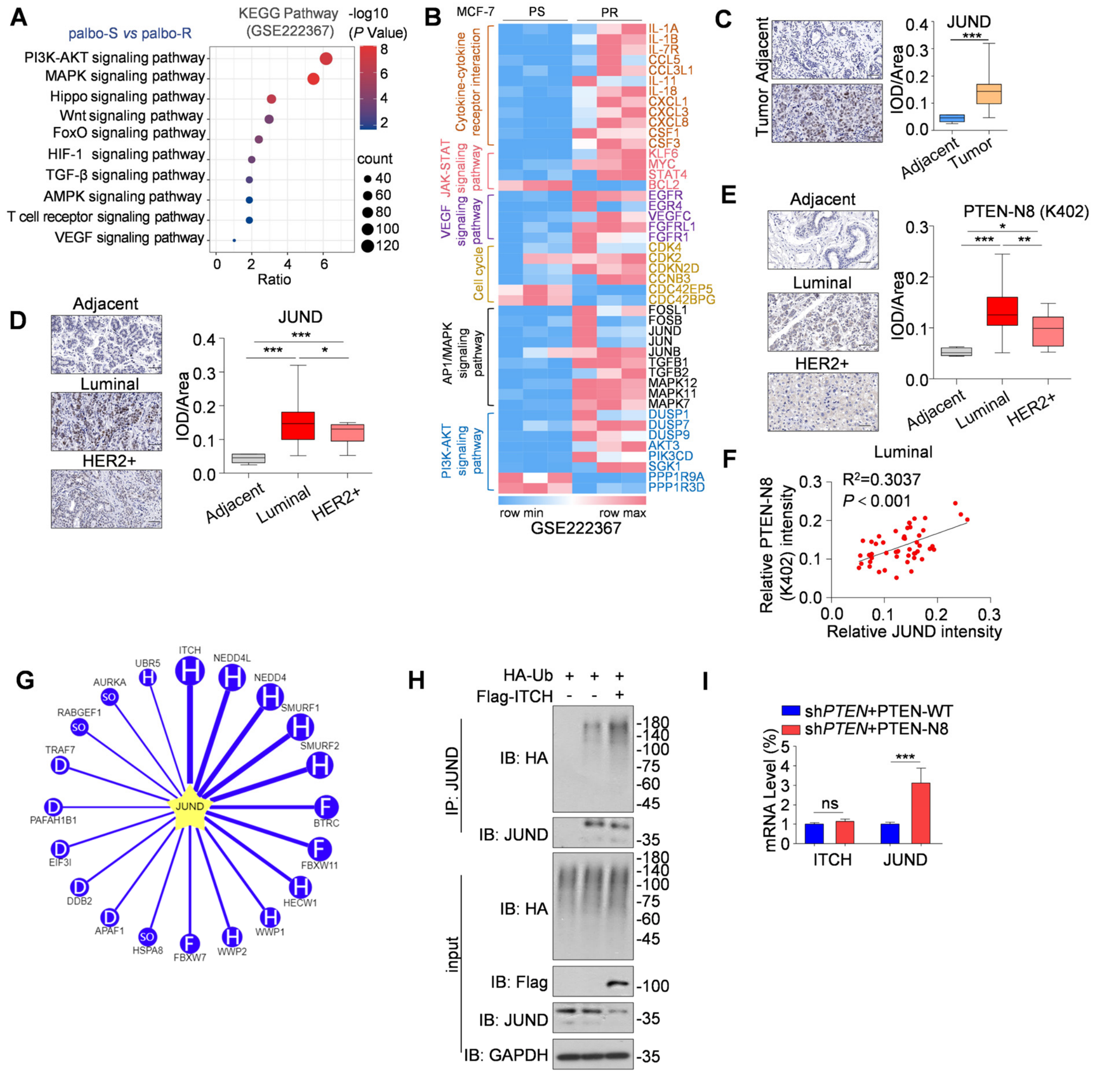

**Supplementary Fig. 2 PTEN neddylation enhances JUND stability in palbociclib-resistant breast cancer.** **A** The bubble charts depict the top ranked pathway analyzed from the KEGG pathway database. Enriched KEGG pathways were identified, which included upregulated or downregulated genes modulated by the palbociclib-resistant MCF-7 cells from the dataset (GSE222367). **B** Heat map of the palbociclib-resistant MCF-7 cells (GSE222367). **C** Representative images from immunohistochemical staining of JUND in breast cancer and matched adjacent tissues. ImageJ was used to perform Semi-quantitative analysis. Data were analyzed using

Student's *t*-test. Scale bars, 50 μm. JUND expression is shown as box plots. **D, E** Representative images from immunohistochemical staining of JUND (**D**) and neddylated PTEN on K402 site (**E**) in different breast cancer subtypes and adjacent tissues are shown. Scale bars, 50 μm. Data were analyzed using the one-way ANOVA test. JUND and PTEN neddylation on K402 expression are shown as box plots. **F** Correlation between JUND and PTEN neddylation on K402 site. **G** Network view of the top 20 predicted E3 ligase proteins of JUND by ubibrowser 2.0 (http: //ubibrowser.bio-it.cn/ubibrowser_v3/). **H** Immunoprecipitation of JUND in HEK293T cells transfected with plasmids of HA-tagged-Ub, Flag-tagged-ITCH and analysis using immunoblot analysis with the indicated antibodies. **I** Analysis of ITCH and JUND mRNA levels in MCF-7 sh*PTEN*+PTEN-WT/Nedd8 cells. *P* values were calculated by Student's *t*-test (**C, I**) and one-way ANOVA test (**E, D**). Error bars, ± S.D. ns, not significant, **P* < 0.05, ***P* < 0.01, ****P* < 0.001.

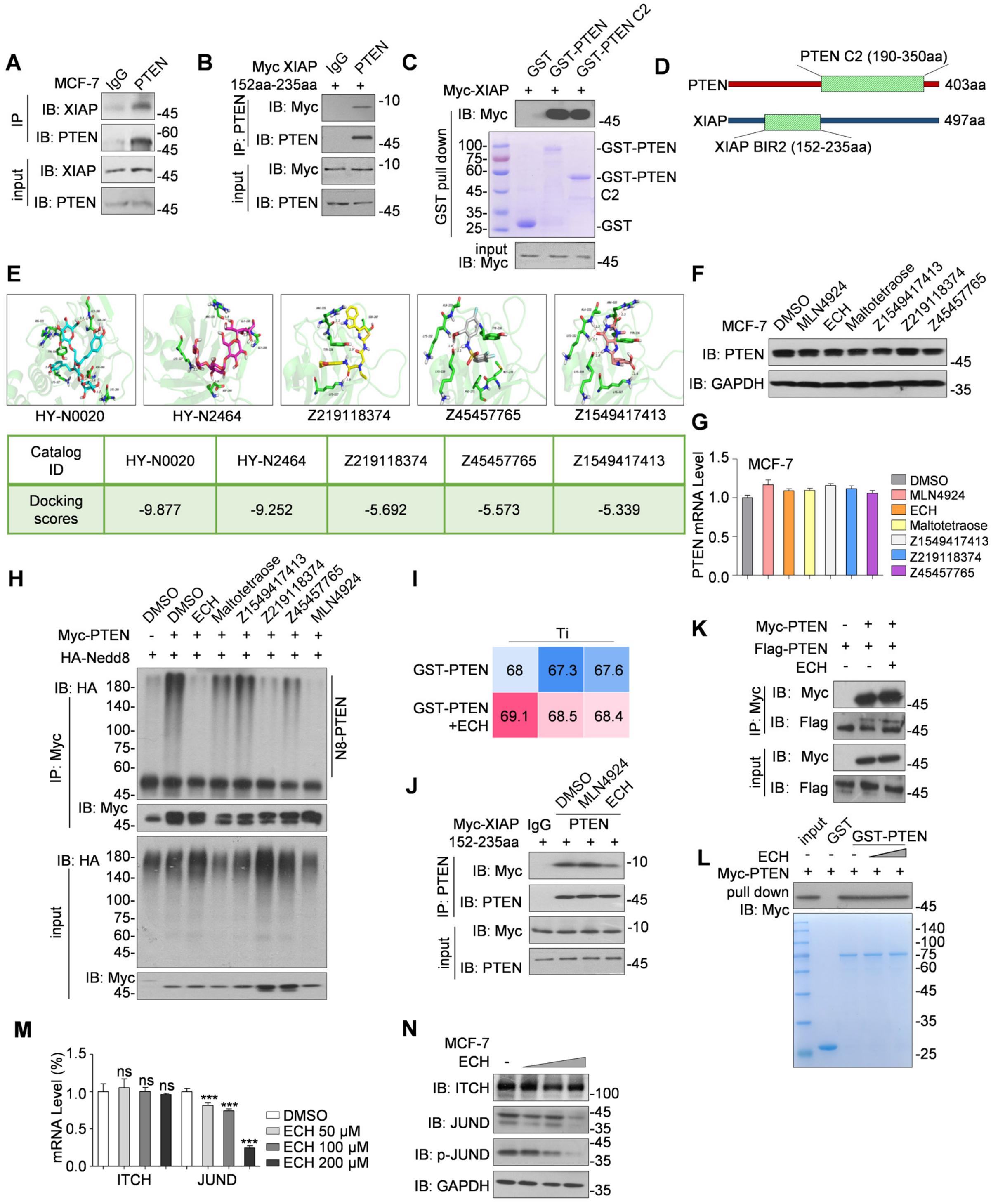


| Catalog ID | HY-N0020 | HY-N2464 | Z219118374 | Z45457765 | Z1549417413 |
|---|---|---|---|---|---|
| Docking scores | -9.877 | -9.252 | -5.692 | -5.573 | -5.339 |

**Supplementary Fig. 3 Echinacoside inhibits the binding of PTEN and XIAP to suppress PTEN neddylation. A** Coimmunoprecipitation of XIAP after immunoprecipitation of the PTEN protein in MCF-7 cells. **B** Immunoprecipitation of PTEN in HEK293T cells transfected with plasmids of Myc-tagged XIAP (152aa-235aa) and analysis using immunoblot with the indicated antibodies. **C** Immunoblot analysis of GST pull-downs or WCL from HEK293T cells transfected with indicated constructs. **D** Diagram illustrating the interacting truncation of PTEN and XIAP. **E** The top five compounds ranked by their molecular docking scores for interaction with both PTEN and XIAP. **F** Immunoblot of WCL from MCF-7 cells treated with indicated drugs. **G** Analysis of PTEN mRNA levels in MCF-7 cells treated with indicate drugs. **H** Immunoblot of anti-Myc immunoprecipitate and whole cell lysate (WCL) from HEK293T cells transfected with indicated constructs. The MCF-7 cells were treated with ECH (200 μM for 24 h), Maltotetraose (50 μM for 24 h), Z1549417413 (50 μM for 24 h), Z219118374 (50 μM for 24 h), Z45457765 (50 μM for 24 h), MLN4924 (1 μM for 12 h) before harvested (**F, G, H**). **I** The inflection temperature (Ti) of GST-PTEN treated with ECH (200 μM for 30 minutes) as determined by TSA assay. **J** Immunoprecipitation of PTEN were obtained from HEK293T cells transfected with Myc-XIAP (amino acids 152-235) and treated with either MLN4924 (0.5 μM for 12 h), Echinacoside (ECH) (200 μM for 24 h). **K** Coimmunoprecipitation of Myc-tagged PTEN after immunoprecipitation of the Flag-tagged PTEN protein in MCF-7 cells. **L** Immunoblot analysis of GST pull-downs or WCL from HEK293T cells transfected with indicated constructs. The concentration of ECH ranged from 50 μM to 200 μM for 24 h. **M** Analysis of ITCH and JUND mRNA levels in MCF-7 cells treated by ECH. The concentration of ECH ranged from 50 μM to 200 μM for 24 h. **N** Immunoblot of WCL from MCF-7 cells for indicated proteins. The concentration of ECH ranged from 50 μM to 200 μM for 24 h. $P$ values were calculated by one-way ANOVA test (**M**). Error bars, ± S.D. ns, not significant, $^{***}P < 0.001$.

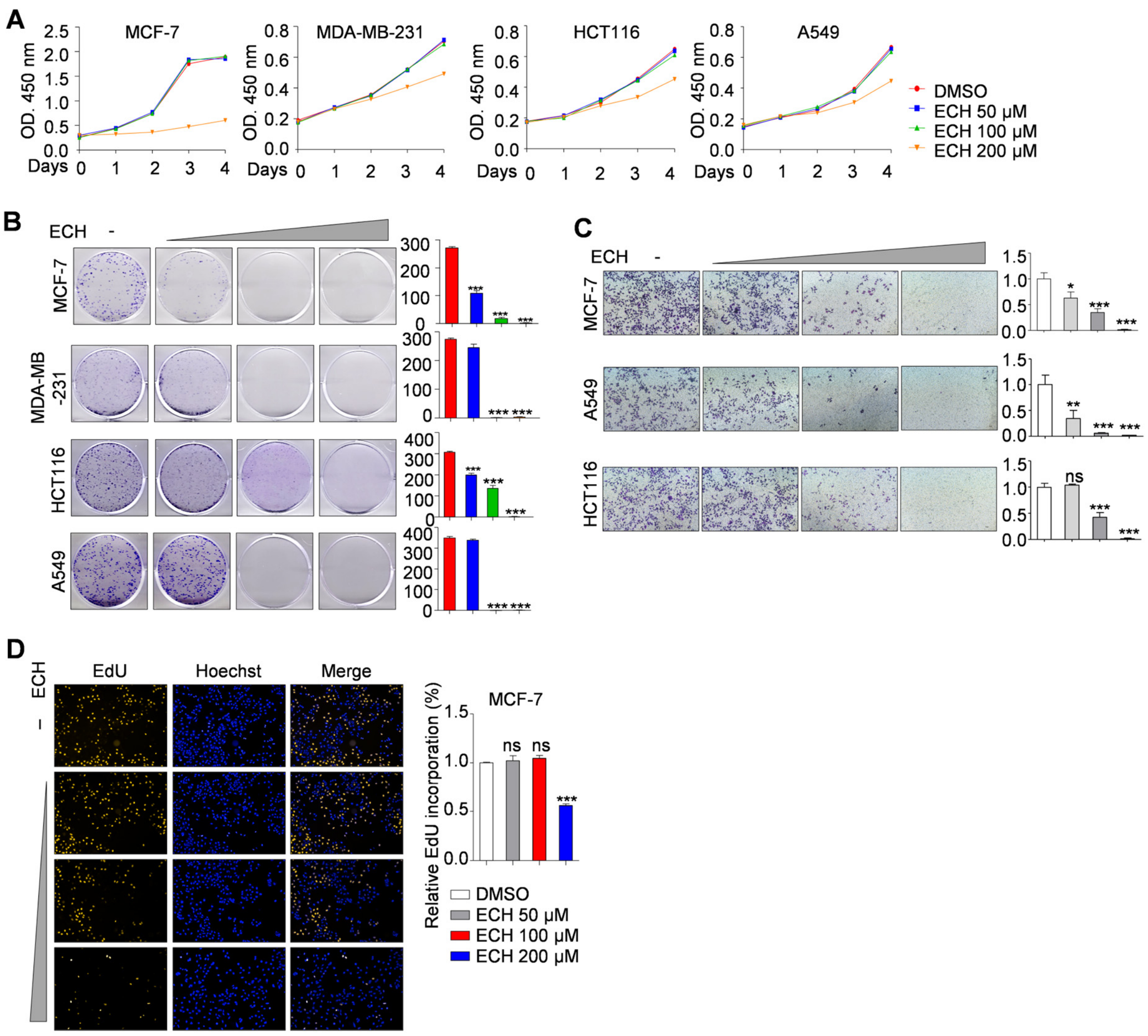


**Supplementary Fig. 4 ECH inhibits the tumor cell invasion and proliferation. A-D** CCK8 assay (**A**), colony formation assay (**B**), cell invasion assay (**C**) and EdU assay (**D**) were performed in the indicated cells treated with ECH. Data are presented as means ± S.D. Results are from a representative experiment performed in triplicate. ImageJ was used to perform quantitative analysis. *P* values were calculated by one-way ANOVA test (**B-D**). Error bars, ± S.D. ns, not significant, **P* < 0.05, ***P* < 0.01, ****P* < 0.001.

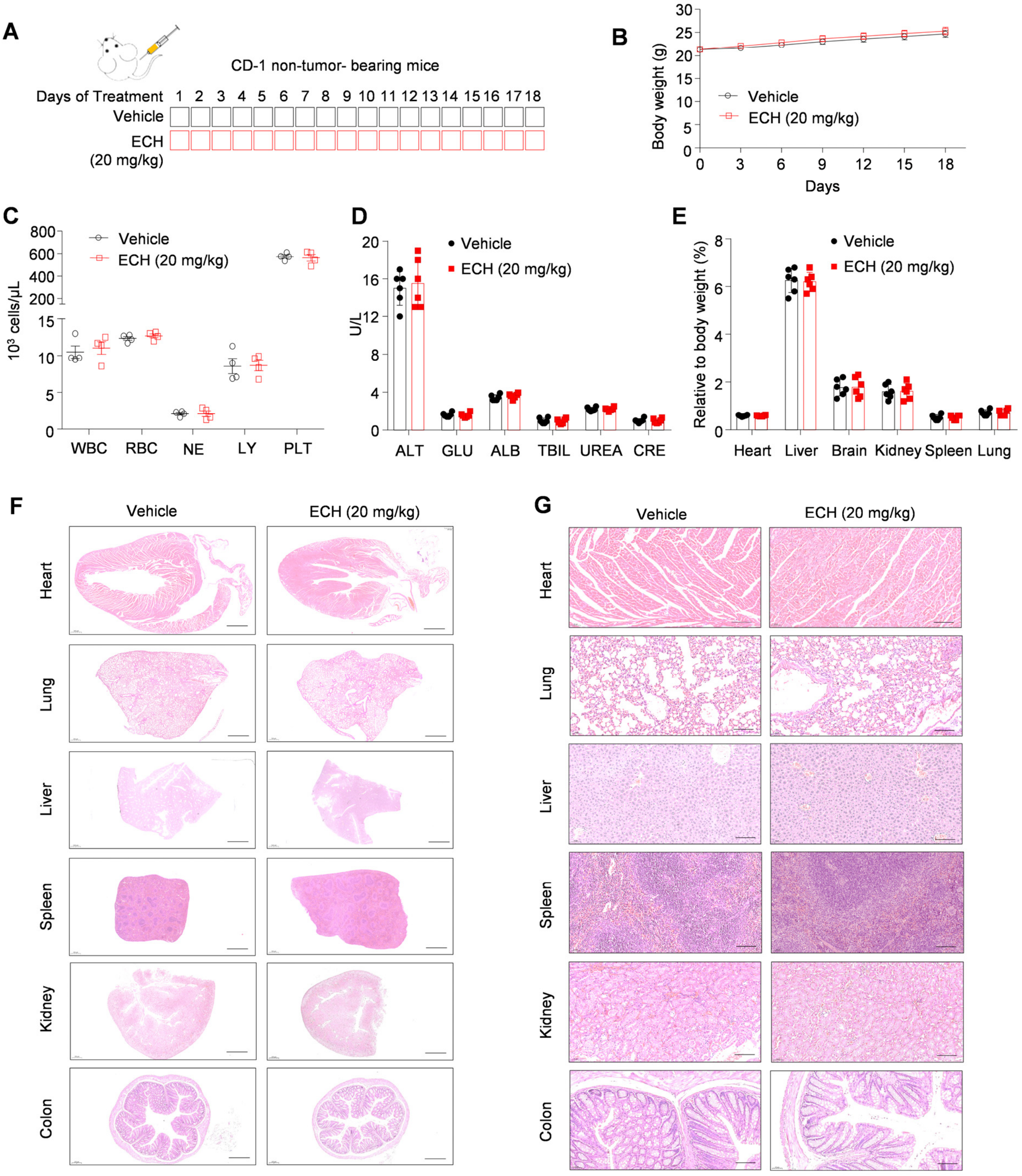


**Supplementary Fig. 5 ECH exhibits low toxicity in CD-1 non-tumor-bearing mice. A** CD-1 non-tumor-bearing mice were treated with saline solution or ECH (20 mg/kg, i.p.) for 18 days. **B** Body weights of CD-1 non-tumor bearing mice were recorded. **C** The blood parameters, including

White Blood Cell count (WBC), Red Blood Cell count (RBC), Neutrophils (NE), Lymphocytes (LY), and Platelet count (PLT) were assessed in CD-1 mice administered either saline solution or ECH. **D** Serum biochemical indicators in CD-1 mice, including ALT, GLU, ALB, TBIL, UREA, CRE. **E** The ratio of organ weights to the overall body weight of the animals was determined. **F, G** H&E staining of heart, lung, liver, spleen, kidney and colon tissue of CD-1 mice. Scale bar, 500 μm (**F**); Scale bar, 50 μm (**G**).

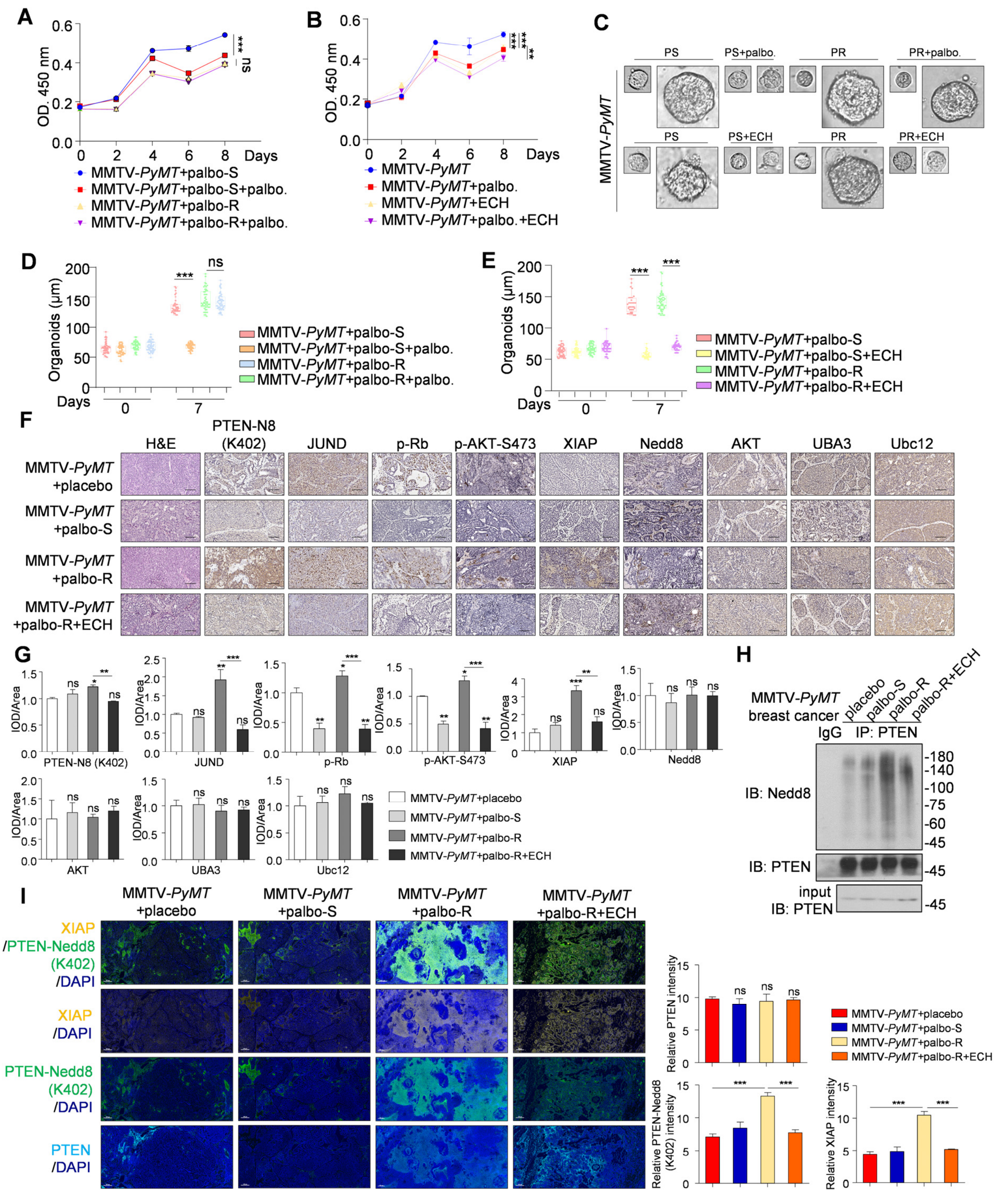

**Supplementary Fig. 6 ECH resists CDK4/6 inhibitor resistance in MMTV-*PyMT* breast cancer mice. A, B** CCK8 assay was performed in palbociclib-resistant breast tumor cells obtained from MMTV-*PyMT* mice (MMTV-*PyMT*+palbo-S and MMTV-*PyMT*+palbo-R). The cells were treated with palbociclib (0.5 μM) for 8 days (**A**) and ECH (200 μM) for 8 days (**B**). **C-E** Breast cancer organoids derived from MMTV-*PyMT* mice (MMTV-*PyMT*+palbo-S and MMTV-*PyMT*+palbo-R) were treated with palbociclib (0.5 μM) and ECH (200 μM) for 7 days respectively. The diameter of organoids was shown as box plots (**D, E**). **F** Representative images from immunohistochemical staining of PTEN neddylation on K402, JUND, p-Rb, p-AKT-S473, XIAP, Nedd8, AKT, UBA3 and Ubc12 in indicated groups. Scale bars, 50 μm. **G** ImageJ was used to perform Semi-quantitative analysis for the expression of indicated protein in (**G**). **H** Immunoblot analysis of anti-PTEN immunoprecipitate and WCL from MMTV-*PyMT* breast cancer tissues to detect PTEN neddylation on K402 site in indicated groups. **I** Representative images from multiplex immunohistochemical staining of neddylated PTEN on K402, PTEN and XIAP in breast cancer tissues of MMTV-*PyMT* mice (n=3). Scale bars, 100 μm. ImageJ was used to perform Semi-quantitative analysis. $P$ values were calculated by one-way ANOVA test (**A, B, D, E, G, I**). Error bars, ± S.D. ns, not significant, *$P < 0.05$, **$P < 0.01$, ***$P < 0.001$.

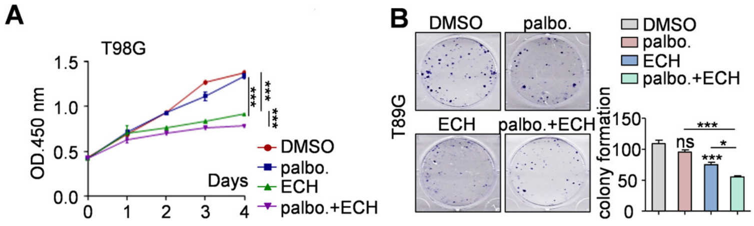


**Supplementary Fig. 7 Combination of ECH enhances the sensitivity of palbociclib in breast cancer.** **A** CCK8 assay was performed in T98G cells treated with ECH (50 μM) and palbociclib (0.5 μM). **B** Colony formation assay was performed in T98G cells treated with ECH (20 μM) and palbociclib (0.1 μM). Data are presented as means ± S.D. Results are from a representative experiment performed in triplicate. ImageJ was used to perform quantitative analysis. *P* values were calculated by one-way ANOVA test (**A, B**). Error bars, ± S.D. ns, not significant, $*P < 0.05$, $**P < 0.01$, $***P < 0.001$.

### SUPPLEMENTARY METHODS

#### RNA-seq Analysis

Total RNA was extracted with TRIzol (Cat# 15596; Invitrogen) and the quality matched the requirement of database sequencing. After that, RNA was digested for reverse transcription, and rRNA was degraded. A complementary DNA library was prepared, and sequencing was performed according to the Illumina standard protocol by Novogene Technology Co., Ltd. (Beijing, China). Reads were normalized and fragments per kilobase per million mapped reads (FPKMs) were calculated through RSEM. MA-plot-based method was used to filter differentially expressed genes (DEGs), and genes with log2 ratio ≥ 2 and *Q*-value ≤ 0.001 were considered significantly regulated.

Gene ontology enrichment and Kyoto Encyclopedia of Genes and Genomes pathway analysis of DEGs were performed using Phyper function in R software. FDR correction was performed on *P*-value, and *Q*-value ≤ 0.05 considered as significant enrichment.

#### Real-time quantitative RT-PCR

MCF-7 cells were harvested and total RNA was extracted using the TRIzol reagent (Invitrogen) according to the manufacturer's instructions. Reverse transcription of 3 μg total RNA was performed by combining 1 μL ReverTraAce-α-™ (Toyobo. No. FSK-100), 4 μL 5 × buffer, 1 μL 10 pmol/μL oligo (dT) primer, 2 μL 10 mM dNTP mix, 1 μL 10 U/μL RNase inhibitor, and water up to a volume of 20 μL. Reaction mixtures were incubated at 30 °C for 10 min, then 42 °C for 20 min, 99 °C for 5 min, 4 °C for 5 min and then diluted with 30 μL of water. Each PCR mixture contained 0.5 μL of cDNA template and primers at a concentration of 100 nM in a final volume of 25 μL of SYBR green reaction mix (Toyobo. No. QPK-201). Standard curves were calculated using cDNA to determine the linear range and PCR efficiency of each primer pair. Reactions were done in triplicate, and relative amounts of cDNA were normalized to GAPDH. The primers used were listed in Supplementary Table 3.

**Thermal shift assay (TSA)**

TychoTM NT.6 checked the protein of GST-PTEN quality. Use 10 μL of sample to find out the quality of GST-PTEN protein in 3 minutes and automatically generates thermal unfolding profiles, identifies inflection temperatures (Ti).

**Cell proliferation assay**

Cells were plated on 96-well plates (2000 cells per well). After adding Cell Counting Kit-8 (Dojindo, CK04) to the wells for 1 h, cell numbers were measured at 450 nm. Each cell line was set up in 3 replicate wells, and the experiment was repeated three times.

**Cell migration assay**

Cell migration assay was performed using Cell culture inserts for 24-well plates with 8.0 μm pores filters, and the filter was pre-coated with 20 μg/ml Fibronectin. A cell suspension in serum-free culture medium ($5 \times 10^4$ cells per well) was added to the inserts, and each insert was placed in the lower chamber containing 10 % FBS culture medium. After 24 h, cells were fixed using 4 % paraformaldehyde for 20 min and stained using 0.1 % crystal violet for 15 min. Cells in upper chamber were carefully removed, and cells migrated through the filter were assessed by photography.

**Colony formation assay**

Cells were seeded into 6-well tissue culture plate with 2000 cells per well. Plates were incubated at 37 °C and 5 % $CO_2$ and colonies were scored 14 days after preparation. Each cell line was set up in 3 replicate wells, and the experiment was repeated three times.

**H&E staining**

Hematoxylin-eosin (H&E) staining was performed to examine breast cancer tissue morphology according to standard protocols. In brief, tissues were fixed with 4 % paraformaldehyde (PFA) overnight. The fixed tissue samples were dehydrated with different concentrations of ethanol and xylol followed by brief washing and staining of cell nuclei with 5 % hematoxylin solution for 10 min. After rinsing in distilled water for 5 min, the stained samples were incubated in 0.1 % HCl-ethanol for 30 s. The samples were then counterstained with eosin solution for 2 min.
