## Supplementary material for "PTEN neddylation aggravates CDK4/6 inhibitor resistance in breast cancer": Supplementary Table 3-clean version.docx

Supplementary Table 3: Primers

| primers for shRNA | |
| --- | --- |
| sh*PTEN* | TGCAGATAATGACAAGGAA |
| sh*NC* | TTCTCCGAACGTGTCACGT |
| primers for qPCR | |
| *PTEN* | TGGATTCGACTTAGACTTGACCT |
|  | TGGATTCGACTTAGACTTGACCT |
| *XIAP* | ACCGTGCGGTGCTTTAGTT |
|  | TGCGTGGCACTATTTTCAAGATA |
| *CDK2* | CCAGGAGTTACTTCTATGCCTGA |
|  | TTCATCCAGGGGAGGTACAAC |
| *JUND* | TCATCATCCAGTCCAACGGG |
|  | TTCTGCTTGTGTAAATCCTCCAG |
| *ITCH* | TGATGATGGCTCCAGATCCAA |
|  | GACTCTCCTATTTTCACCAGCTC |
| *GAPDH* | GGAGCGAGATCCCTCCAAAAT |
|  | GGCTGTTGTCATACTTCTCATGG |
| primers for ChIP-qPCR | |
| IL-1β promoter | CAGAGTTCCCCAACTGGTACATC |
|  | GGGAAGGCATTAGAAACAGTGTC |
| IL-6 promoter | TCTGCAAGATGCCACAAGGT |
|  | TGAAGCCCACTTGGTTCAGG |
| IL-17 promoter | GCCTTTGTGATTGTTTCTTGCAG |
|  | CCTTGCCCAAAGAAACCCTCTC |
| CSF2 promoter | TGTCGGTTCTTGGAAAGGTTCA |
|  | TGTGGAATCTCCTGGCCCTTA |
| MMP13 promoter | CAACCATGGGGCTCAATCCT |
|  | CTTACGTGGCGACTTTTTCTTTTC |
| primers for routine genotyping | |
| MMTV-*PyMT* (TG) | GGAAGCAAGTACTTCACAAGGG |
|  | GGAAAGTCACTAGGAGCAGGG |
|  | CAAATGTTGCTTGTCTGGTG |
|  | GTCAGTCGAGTGCACAGTTT |
| primers for constructs and mutagenesis | |
| Myc-XIAP (152-235aa) | CCGGAATTCATGGGTTTCTTTATACTGGTGA |
|  | CCGCTCGAGTTACTGAGTCTCCATATTGCCATC |
| Myc-XIAP (152-235aa) Y154F | GGAATTCCGATGACCATATTTCCGAGGAAC CCTGCCATGTATAGTGAAGAAGCT |
|  | CCGCTCGAGTTAAAGATTCCGGCCCAAAAC AAAGAAGCAATTAGGAAAGTG |
| Myc-XIAP (152-235aa) N234A | CCGGAATTCATGGAAGAAGCTAGATTAAAG TCCTTT |
|  | CCGCTCGAGTTAAAGTGCCCGGCCCAAAAC AAAGAA |
| GST-PTEN C2 | AAGAATTCCCAGTGGCACTGTTG |
|  | CTCGAGTCATGTTTTTGTGAAGTACAG |
