## Supplementary material for "PTEN neddylation aggravates CDK4/6 inhibitor resistance in breast cancer": Supplementary Table 4-clean version.docx

Supplementary Table 4: Sequences for RNA interference

| Region | 5' to 3' |
| --- | --- |
| *ITCH-1* | GGAUCACAACUUGGUUCAATT |
| *ITCH-2* | GGAAAUGUACUUCUCCGUUTT |
| Negative control | UUCUCCGAACGUGUCACGUTT |
